# Condensin and topoisomerases cooperate to relieve topological stress at stalled replication forks

**DOI:** 10.1101/2025.08.06.668895

**Authors:** Mégane Da Mota, Axel Delamarre, Antoine Barthe, Jessica Jackson, Alba Torán-Vilarrubias, Cyril Ribeyre, Alessandro Vindigni, Yea-Lih Lin, Philippe Pasero, Armelle Lengronne

**Affiliations:** Institut de Génétique Humaine, Univ Montpellier, CNRS, Montpellier, France; Division of Oncology, Department of Medicine, Washington University School of Medicine, St. Louis, MO 63110, USA; Institut de Génétique Moléculaire de Montpellier, Univ Montpellier, CNRS, Montpellier, France

**Author notes:** Laboratoire d’Eco-Anthropologie, CNRS, Muséum National d’Histoire Naturelle, Université Paris Cité, Paris, France.

## Abstract

Resolving complex topological structures at replication forks is vital for successful DNA replication, but the mechanisms are little understood. Evidence from diverse eukaryotes suggests that condensin – which promotes chromosome condensation in M phase – might also act during S phase to facilitate relaxation of torsional stress by topoisomerases. Here, we show in yeast and human cells that condensin binds stressed replication forks, where it cooperates with topoisomerases I and II to promote resection of the nascent DNA and restart replication. Our findings suggest that condensin acts with topoisomerase I at reversed forks to convert positively supercoiled DNA into structures that are subsequently relaxed by topoisomerase 2, allowing the fork to resume replication. These findings uncover an important, evolutionarily conserved role for condensin in handling topological constraints at arrested forks that is reminiscent of its function in chromosome segregation and might prevent formation of toxic chromosome structures during fork arrest and reversal.

## INTRODUCTION

Unresolved or persistently stalled replication forks are vulnerable structures susceptible to nucleolytic attacks and breakage, ultimately driving genome instability. The cell has several mechanisms to overcome fork stalling, including replication fork reversal^1–3^. Replication fork reversal remodels the three-way junction of the replication fork into a four-way junction by reannealing the parental strands and annealing the newly synthesized (nascent) DNA strands to each other to form a fourth strand. This process is mediated by the RAD51 recombinase and DNA translocases such as SMARCAL1^4^. The four-way junction that forms upon fork reversal is a transient structure that slows replication fork progression and facilitates the resumption of replication^3^.

Fork reversal is directly influenced by the topological context in which it takes place. This process is initially driven by the positive torsional stress accumulating in front of the fork during DNA replication^5^. However, further extension of the reversed arm generates positive supercoiling along sister chromatids, limiting the extent of fork reversal^6^ and is relieved by DNA topoisomerase 2 (TOP2)^7^. TOP2 is also important in M phase when it decatenates replicated sister chromatids to allow chromosome segregation^8^. Condensin, a structural maintenance of chromosomes (SMC) protein complex that drives chromosome condensation in M phase, favors TOP2 decatenation activity^9^. This regulatory function of condensin might also be important in interphase.

Two different condensin complexes promote chromosome condensation and segregation in vertebrates^9^. Condensin II localizes to the nucleus during interphase, associates with chromatin and acts from S phase to prophase to resolve sister chromatid intertwines^10^. By contrast, condensin I binds chromosomes in prometaphase, only after nuclear envelope breakdown^9^. Moreover, in the budding yeast *S. cerevisiae*, the unique condensin complex is required for chromosome segregation, and its association with chromatin throughout the cell cycle suggests a role outside of mitosis^11–16^.

Numerous studies suggest that condensin begins to act in S phase and may play a role in preventing DNA damage. In various organisms, mutations in or depletion of condensin result in increased sensitivity to DNA damage and replication stress. In *Arabidopsis thaliana*, condensin II mutants are highly sensitive to DNA-damaging agents that cause double-strand breaks and replication blocks^17^. Exposure of condensin II-depleted cells to aphidicolin leads to severe chromosomal disintegration, revealing the importance of condensin in maintaining chromosome architecture under replication stress^18^. In mouse embryonic stem cells, depletion of the condensin subunit Smc2 results in elevated levels of γ-H2AX foci^19^, indicative of increased DNA damage during S and G_2_ phases. A similar effect is observed in condensin II missense mutants^20^. In the fission yeast *S. pombe*, condensin prevents transcription-induced DNA damage in G_2_-arrested cells^21^. Other condensin mutations make cells hypersensitive to hydroxyurea, which slows DNA replication, and to the DNA topoisomerase inhibitor camptothecin, further highlighting the importance of condensin in managing replication stress^22^. Altogether, these findings suggest that condensin might facilitate relaxation of topological constraints during S phase, particularly during replication stress.

Here, we investigated whether condensin and topoisomerases cooperate to relieve torsional stress in S phase due to replication stress, as they do in mitosis. We show that condensin is recruited to stressed replication forks in budding yeast where it cooperates with Top1 and Top2 to promote fork restart. In human cells, condensin II specifically promotes fork processing and restart under conditions of replication stress. Condensin II acts with TOP1 and TOP2A to promote fork slowing and resection, consistent with a model in which condensin II and TOP1 act at stalled forks to convert positively supercoiled DNA into structures that are subsequently relaxed by TOP2A, thus avoiding formation of DNA knots and intertwined sister chromatids that might result in genomic instability.

## RESULTS

### Condensin is recruited to stressed replication forks in an MRX-dependent manner

To determine whether yeast condensin is required for growth under replication stress conditions, we compared the growth of *S. cerevisiae* cells carrying the wild-type allele of the condensin core subunit Smc2, *SMC2*, with that of those carrying the thermosensitive (ts) *smc2-8* allele, in the presence of MMS or HU. MMS impedes replication fork progression by alkylating DNA, whereas HU slows DNA synthesis by depleting dNTP pools and causing oxidative stress. At 30°C, a semi-permissive temperature for the *smc2-8* allele, both strains grew at the same rate in the absence of drugs. The growth of *smc2-8* cells was significantly impaired, however, by chronic exposure either to 100 mM HU or to 0.033% MMS (**Fig. 1a**). A similar growth defect on MMS-containing medium was observed when the other condensin core subunit, Smc4 tagged with an auxin-inducible degron (AID) was partially depleted by adding a low dose of the auxin IAA (**Supplementary** Fig. 1a).

**Fig. 1.**
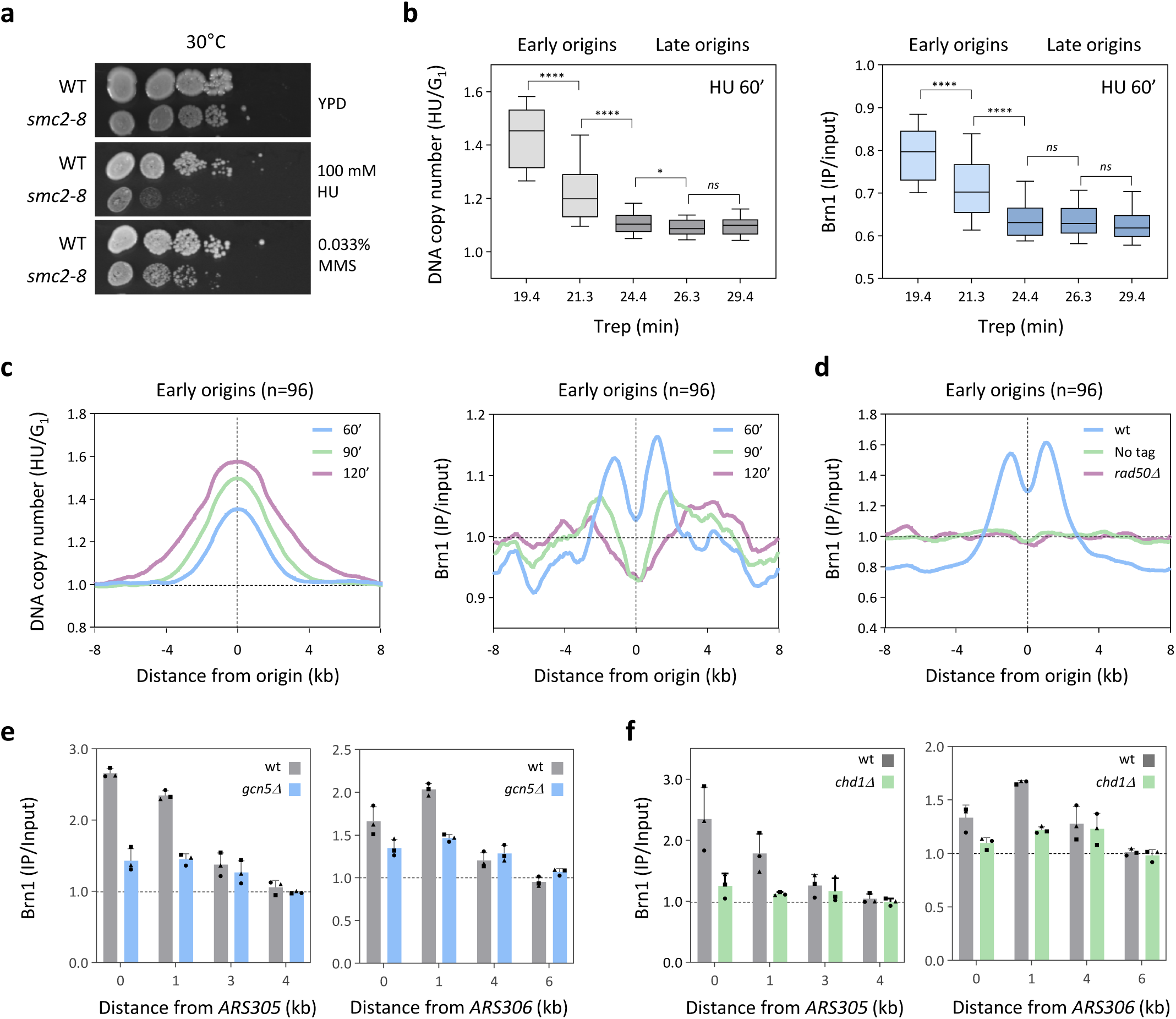
Condensin is recruited to stressed replication forks in budding yeast. **a** Condensin promotes cell growth under replication stress conditions. Wild type (WT) and thermosensitive *smc2-8* cells were grown for 4 days on YPD medium containing 0.033% MMS or 100 mM HU at the semi-restrictive temperature of 30°C. **b** Condensin is enriched at early origins in HU-treated cells. Cells expressing a tagged condensin subunit (*BRN1-PK_9_*) were arrested in G_1_ with α-factor and were released for 60 minutes into medium containing 200 mM HU. Brn1 was immunoprecipitated and changes in DNA content (left) and Brn1 enrichment (right) was determined by ChIP-seq at all annotated *S. cerevisiae* replication origins as described^27,71^. DNA copy number and Brn1 enrichment were determined for 2 kb intervals and are expressed for groups of early and late origins sorted according to their replication time (Trep,^25^). Mean Trep is indicated for each bin. Box and whiskers indicate 25^th^–75^th^ and 10^th^–90^th^ percentiles, respectively. **** p < 0.0001; * p<0.1; *ns* not significant; Mann-Whitney rank sum test. **c** Condensin follows replication forks in HU-treated cells. Cells were released synchronously into S phase for 60, 90 and 120 minutes in the presence of 200 mM HU and the distribution of DNA content (left) and Brn1 enrichment (right) was determined by ChIP-seq for 16 kb intervals centered on 96 early origins. **d** Condensin is recruited to stressed forks by the MRX complex. Brn1 enrichment at 96 early origins was quantified by ChIP-seq in wild type (untagged), *BRN1-PK_9_* and *rad50∆ BRN1-PK_9_* cells at 60 minutes post G_1_ release into 200 mM HU. **e**, **f** Condensin binding to HU-arrested forks depends on the Gcn5 histone acetyl transferase and on the Chd1 chromatin remodeler. Brn1 enrichment at increasing distances from the early origins *ARS305* and *ARS306* was determined by ChIP-qPCR in wild type (grey), *gnc5∆* (blue) and *chd1∆* (green) cells. Cells were released from G_1_ into S phase for 60 minutes in the presence of 200 mM HU. Brn1 enrichment was normalized to unreplicated loci as described^27^. Mean and SD correspond to three independent experiments.

Condensin binds to multiple sites in the yeast genome throughout the cell cycle, including replication origins^23^. To characterize further the interaction of condensin with replication sites, we analyzed the distribution of the condensin subunit Brn1 by ChIP-seq in cells synchronized in S phase by release from G_1_ into S phase for 60 min in the presence of 200 mM HU to arrest forks 2-3 kb from early origins^24^. This analysis showed that condensin is absent from replication sites in G_1_-arrested cells and is thus specific to the S phase (**Supplementary** Fig. 1b). To determine whether this binding depends on origin firing, we analyzed origin activation in HU-arrested cells (t=60 min) compared with G_1_-arrested cells. Origin firing induces a local increase in DNA copy number, expressed as the ratio of genomic DNA reads (HU/G_1_) for 2-kb intervals centered on annotated yeast origins (n=385). This ratio was plotted against the replication time (Trep;^25^) for bins of 77 origins (**Fig. 1b**). Under these experimental conditions, early origins fired (HU/G_1_ ratio > 1), whereas late origins, normally activated 24 min after release from G_1_ phase, were repressed by the replication checkpoint, consistent with earlier studies^24,26^. We then calculated Brn1 enrichment at the same intervals, and found significant enrichment at early origins but not at late origins (**Fig. 1b**). Together, these data suggest that condensin is recruited to replication sites after origin firing in HU-treated cells.

To investigate whether the binding of condensin to these replication sites changes over time under conditions of replication stress, we analyzed the distribution of Brn1 by ChIP-seq in cells released synchronously from G_1_ into S phase for 60, 90 and 120 min in the presence of 200 mM HU. We reported previously that forks progress at a rate of 0.1 kb/min under these conditions^24^. To determine whether condensin complexes progress with replication forks in HU-treated cells, we plotted the variation of DNA copy number and Brn1 enrichment over time for 16-kb intervals centered on 96 early origins (**Fig. 1c**). Over time, condensin moved away from early origins, as it is the case for changes in DNA copy number (**Fig. 1c**). Together, these data suggest that the condensin complex is recruited to early replication origins during HU-mediated replication stress and travels with replication forks.

The Mre11–Rad50–Xrs2 (MRX) complex acts together with histone modifiers and nucleosome remodelers Gcn5 and Chd1 to increase chromatin accessibility and promote loading of another SMC family protein complex, the sister chromatid cohesion protein cohesin, at stressed forks^27^. To investigate whether condensin recruitment to stressed forks also involves the MRX complex, we synchronized wild-type and *rad50Δ* cells in S phase by using HU as above, and determined Brn1 enrichment at early origins by ChIP-seq (**Fig. 1d** and **Supplementary** Fig. 1b). The amount of Brn1 at early origins in *rad50Δ* cells was reduced to the same level as in the control cells lacking the Pk tag. We confirmed this finding specifically at the early origins *ARS305* and *ARS306* (**Supplementary** Fig. 1c) by using ChIP-qPCR as described previously^27^. We also measured Brn1 levels at *ARS305* and *ARS306* by ChIP–qPCR in cells lacking the histone acetyltransferase Gcn5 (*gcn5Δ*) and the chromatin remodeler Chd1 (*chd1Δ*). In wild-type cells, Brn1 was located behind HU-arrested forks, which progress by 2– 3 kb from the origin under these conditions^24^; by contrast, no Brn1 enrichment was detected in *gcn5Δ* and *chd1Δ* mutants (**Fig. 1e** and **1f**). Overall, these data indicate that condensin is recruited to replication stress sites through a mechanism involving the MRX complex and chromatin remodeling, reminiscent of the recruitment of cohesin^27^.

### Topoisomerases cooperate with condensin to promote fork restart

The fission yeast condensin mutant *cnd2-1* is sensitive to genotoxic agents, likely because it is unable to activate the DNA replication checkpoint^28^. To determine whether this checkpoint is active in the absence of condensin in budding yeast, we used the repression of late origins as a readout for its timely activation^26^ in pMET3-*YCG1*-AID cells, in which expression of the condensin subunit Ycg1 can be suppressed by growth in rich medium containing methionine and degraded by the addition of auxin, as previously described^29^. Wild-type and pMET3-*YCG1*-AID cells were first grown in synthetic medium lacking methionine to maintain *YCG1* expression, and arrested in G_1_ phase with α-factor; then they were placed in rich medium containing methionine and auxin^29^ to induce *YCG1* transcription shutoff and Ycg1 degradation (**Supplementary** Fig. 2a). The cells were then released from G_1_ phase arrest and allowed to enter S phase for 60 min in the presence of 200 mM HU, and changes in DNA copy number at early and late origins were monitored by qPCR, as reported previously^26^. This analysis showed that late origins were properly repressed in condensin-depleted cells as in wild-type cells (**Supplementary** Fig. 2b), indicating that condensin is dispensable for checkpoint activation at stressed forks.

To investigate whether condensin promotes fork progression under conditions of replication stress, we monitored the impact of HU on DNA synthesis along individual DNA fibers by using the DNA combing technique, as described previously^30^. Wild-type and *smc2-8* mutant cells were arrested in G_1_ phase and released synchronously into S phase for 90 or 180 min at the restrictive temperature of 35°C in the presence of the synthetic nucleoside analogue bromodeoxyuridine (BrdU, which incorporates into DNA in place of thymidine and can be detected fluorescently) and 200 mM HU, followed by DNA combing analysis. In control cells, the length of BrdU tracks increased by 8.7 kb between 90 and 180 min in HU-containing medium, whereas in *smc2-8* mutants this increase was reduced 2.8-fold (to 3.1 kb) when compared with control cells (**Fig. 2a** and **Supplementary Fig.2c**). A significant reduction in BrdU track length was also detected in Ycg1-depleted cells (**Supplementary** Fig. 2d), even though condensin is dispensable for normal fork progression in the absence of HU (**Supplementary** Fig. 2e). Together, these data indicate that condensin is required for optimal fork progression under conditions of replication stress.

**Fig. 2.**
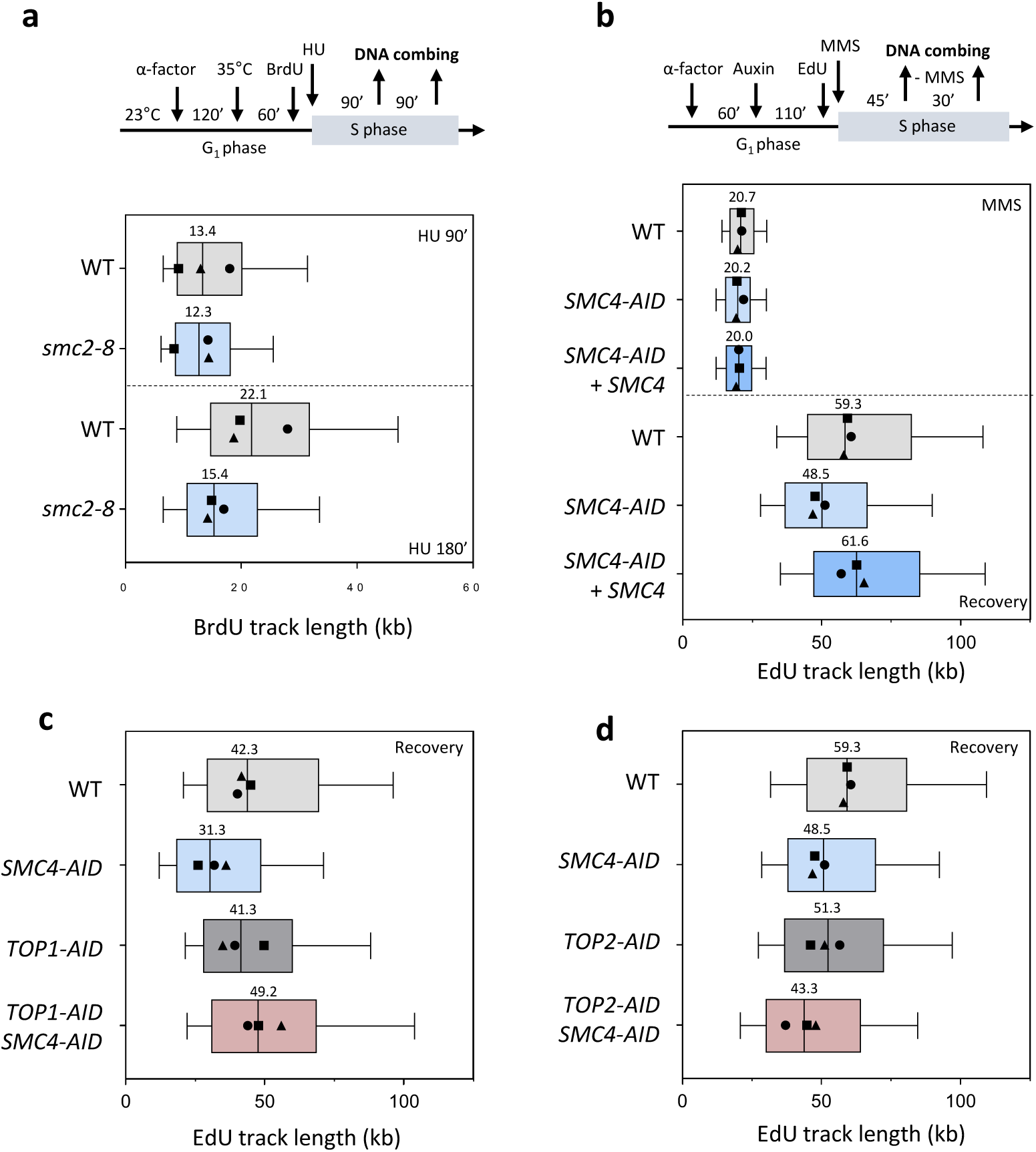
Condensin promotes fork restart after replication stress in budding yeast. **a** Condensin promotes fork progression in the presence of HU. Wild type and *smc2-8* cells were synchronized in G_1_ with α-factor, shifted at 35°C for 60 minutes and released into S phase in the presence of 200 mM HU. Newly replicated DNA was labeled with BrdU for 90 and 180 minutes and fork progression was monitored by DNA combing. The distribution of BrdU track length is shown for three independent experiments. Box and whiskers indicate 25^th^–75^th^ and 10^th^–90^th^ percentiles, respectively. Median length is indicated. **b** Condensin is required for timely fork restart after MMS exposure. Wild type, *SMC4*-PK_3_-AID and *SMC4*-PK_3_-AID + *SMC4*-HA_3_ cells were synchronized in G_1_ with α-factor and released into S phase in the presence of 0.05% MMS. Auxin was added 60 minutes before release from the G_1_ arrest. Cells were released for 45 minutes into S phase in the presence of MMS and EdU, then MMS was removed to let cells recover for 30 minutes. The length of EdU tracks was determined after 45 minutes in MMS and after recovery. The distribution of EdU tracks length is shown for three independent experiments. Box and whiskers indicate 25th–75th and 10th–90th percentiles, respectively. **c** Top1 depletion restores fork restart in the absence of condensin. Fork restart after MMS exposure was monitored by DNA combing as indicated in panel b in WT, *SMC4*-PK_3_-AID, *TOP1*-AID and *SMC4*-PK_3_-AID, *TOP1*-AID cells. The distribution of EdU tracks length is shown for three independent experiments. **d** Condensin acts with Top2 to promote fork restart after MMS exposure. Fork restart was monitored by DNA combing in wild-type, *SMC4*-PK_3_-AID, *TOP2*-MYC-AID and *SMC4*-PK_3_-AID, *TOP2*-MYC-AID cells as indicated in panel b. The distribution of EdU track length is shown for three independent experiments.

Since forks undergo multiple rounds of arrest and restart when dNTP concentrations are low, the slow fork progression measured in HU-treated *smc2-8* mutants might reflect a role for condensin in fork restart. To address this possibility, cells were exposed to 0.05% MMS to stall forks by alkylating DNA, and the length of BrdU tracks was measured by DNA combing after removal of MMS. More specifically, wild-type and *SMC4*-AID cells were arrested in G_1_ phase with α-factor and, 1 h after α-factor addition, auxin was added to degrade Smc4. The α-factor was then removed and the cells were allowed to enter S phase for 45 min in the presence of the thymidine analogue 5-ethynyl-2′-deoxyuridine (EdU) and 0.05% MMS to induce acute fork arrest. DNA combing analysis revealed that this treatment impeded DNA synthesis to a similar extent in Smc4-depleted cells as in wild-type cells (**Fig. 2b** and **Supplementary** Fig. 2f). The MMS was then removed and the length of replicated DNA tracks was measured 30 minutes later by EdU incorporation. This analysis showed that EdU tracks were 27% shorter in Smc4-depleted cells after MMS removal than in wild-type cells and in Smc4-depleted cells complemented by ectopic expression of wild-type Smc4 (**Fig. 2b** and **Supplementary** Fig. 2f), indicating that condensin is required for efficient fork restart.

Several lines of evidence suggest that the role of condensin at stalled replication forks might be linked to the activity of DNA topoisomerases. First, condensin functions are closely linked to the activity of topoisomerases I and II. Moreover, condensin-mediated DNA overwinding may be crucial for chromosome decatenation by topoisomerase II^31–34^. Also, topoisomerases play a crucial role in the progression of replication forks by relieving positive superhelical tension^35^. To investigate the interaction of condensin with topoisomerases at stalled replication forks, we examined the ability of cells to restart MMS-arrested replication forks when depleted of topoisomerase I or topoisomerase II, or also depleted of condensin. Wild-type, *SMC4*-AID, *TOP2*-AID, *TOP1*-AID, *SMC4*-AID *TOP2*-AID, and *SMC4*-AID *TOP1*-AID cells were arrested in G_1_ phase by addition of α-factor, and auxin was added 1 h later to induce degradation of the AID-tagged proteins (**Supplementary** Fig. 2i and 2j). The α-factor was removed to allow the cells to progress into S phase in the presence of EdU and 0.05% MMS, and the length of EdU tracks was measured by DNA combing before or after release from the MMS arrest. Whereas no significant difference in EdU track length during the MMS treatment was observed in any of the depleted cells when compared with wild-type cells (**Supplementary** Fig. 2g and 2h), the tracks measured after recovery from MMS were shorter in *SMC4*-AID and *TOP2*-AID cells, but not in *TOP1*-AID cells (**Fig. 2c, 2d, Supplementary** Fig. 2g and 2h). Simultaneous depletion of Smc4 and Top1 restored fork restart in Smc4-depleted cells, suggesting that condensin is required for fork restart in the presence of Top1 but not in its absence (**Fig. 2c** and **Supplementary** Fig. 2g). By contrast, simultaneous depletion of Smc4 and Top2 had no additional effect on fork recovery when compared with depletion of these proteins individually, indicating that condensin and Top2 may function in a common pathway required for efficient fork restart following replication stress (**Fig. 2d** and **Supplementary** Fig. 2h). Together, these data reveal a complex interplay between condensin, Top1 and Top2 at stalled forks that is important to promote fork restart.

### Condensin II travels with replication forks in human cells and promotes fork restart

To determine whether condensin also acts at replication forks in human cells, we labeled HeLa S3 cells for 20 min with 10 µM EdU under normal growth conditions and analyzed the proteins associated with nascent DNA by using iPOND (isolation of proteins on nascent DNA) coupled with mass spectrometry (iPOND-MS)^36^. Smc2 and Smc4, two SMC subunits common to condensin I and II complexes, were associated with nascent DNA (**Fig. 3a**), consistent with previous reports^18,36^, and the amounts decreased upon thymidine chase, as did the amount of PCNA, used here as a positive control; by contrast, the amount of histone H3 did not change. These data suggest that condensin travels with replication forks.

**Fig. 3.**
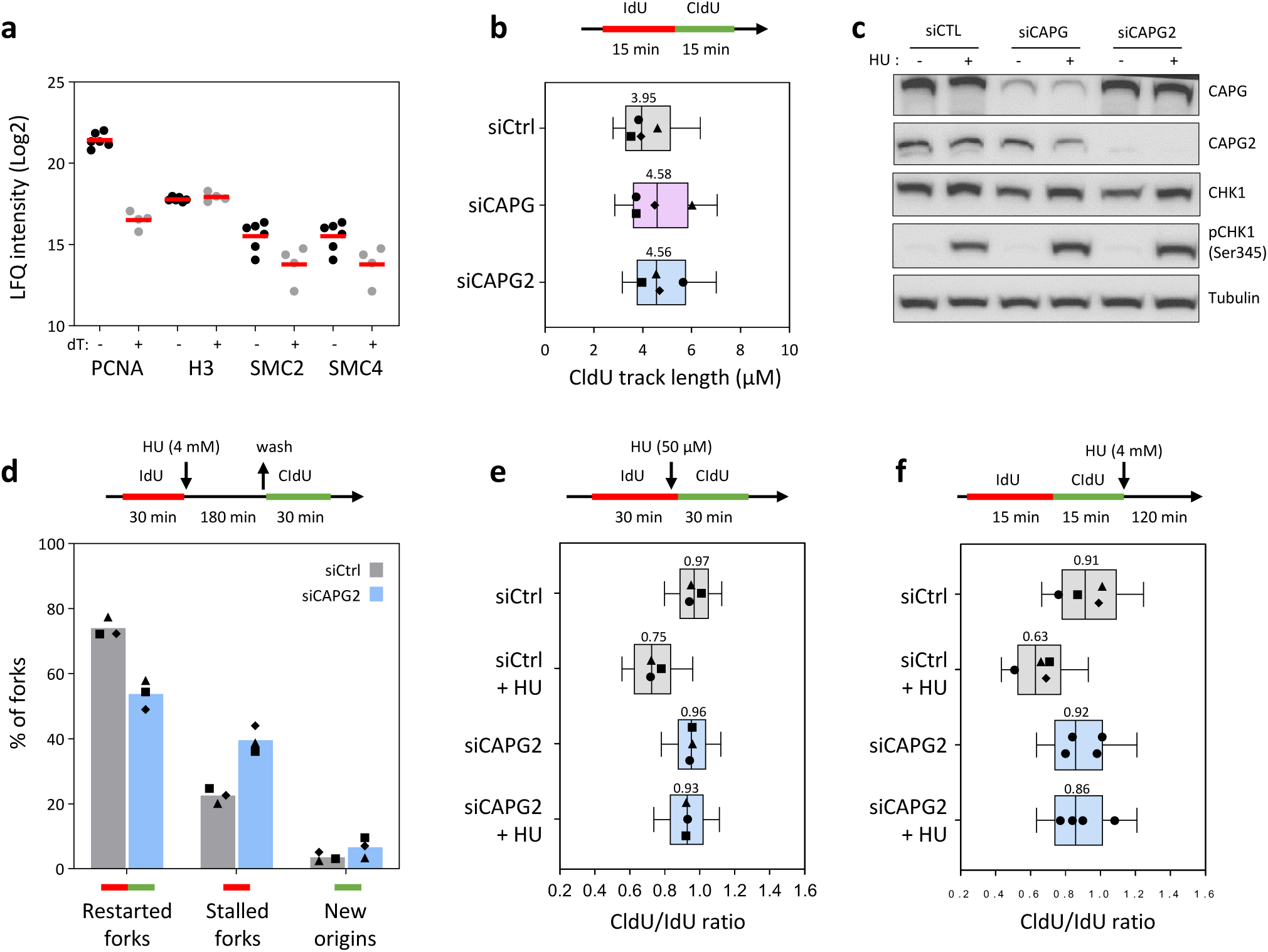
Condensin II promotes fork processing and restart in human cells. **a** Condensin binds nascent DNA. HeLa-S3 cells were labelled for 20 minutes with 10 μM EdU and chased for 60 minutes with thymidine. Proteins associated to nascent DNA before and after the thymidine chase were analyzed by iPOND-MS as described^77^. **b** Condensin I and II are dispensable for normal fork progression. HeLa-S3 cells depleted for condensin I (siCAPG) or condensin II (siCAPG2) for 48 hours with siRNAs were sequentially labeled with IdU and CldU for 15 min before DNA fiber spreading. The length distribution of CldU tracks length is shown for four independent experiments. Box and whiskers indicate 25^th^-75^th^ and 10^th^-90^th^ percentiles, respectively. Median length is indicated. **c** The ATR-CHK1 pathway is functional in the absence of condensin II. HeLa-S3 cells were transfected with siCtrl, siCAPG and siCAPG2 for 48 hours. Cells were then treated with 4 mM HU for 2 hours and the activation of CHK1 was detected with an anti-pCHK1 (S345) antibody. CAPG and CAPG2 depletion was verified by Western blotting. Total CHK1 and tubulin were used as loading controls. **d** Condensin II promotes fork restart. HeLa-S3 cells were transfected with siCtrl or siCAPG2 for 48 hours and were treated with 4 mM HU for 3 hours after a 30 minutes IdU pulse. IdU and CldU tracks were analyzed by DNA fiber spreading 30 minutes after HU removal and CldU addition. Red and green signals are indicative of fork restart. Red only tracks correspond to stalled forks and green tracks to new origin firing (n = 3). **e** Condensin II is required for fork slowing after exposure to a low dose of HU. U2OS cells were transfected with siCtrl and siCAPG2 for 48 hours. Cells were first labelled for 30 minutes with IdU, and CldU was then added for 30 minutes in the presence of 50 μM HU. The length of IdU and CldU tracks was determined by DNA fiber spreading and was plotted as the ratio of CldU to IdU (n=3). Box and whiskers indicate median, 25th–75th and 10th–90th percentiles. **f** Condensin II promotes the resection of nascent DNA at HU-arrested forks. HeLa-S3 cells were transfected with siCtrl and siCAPG2 for 48 hours and were sequentially labeled for 15 minutes with IdU and CldU. Cells were either collected immediately or treated for 2 hours with 4 mM HU before DNA fiber analysis. The ratio of CldU to IdU track length was plotted for 4 independent experiments. Box and whiskers indicate median, 25th–75th and 10th–90th percentiles.

To determine whether condensin participates in DNA synthesis, we depleted the condensin I subunit CAPG and the condensin II subunit CAPG2 from HeLa S3 cells by using siRNAs and monitored the effect of depletion on replication fork progression. Exponentially growing cells were sequentially pulse-labeled with the thymidine analogues iododeoxyuridine (IdU) and chlorodeoxyuridine (CldU) for 15 min each. The DNA molecules were stretched on glass slides by DNA fiber spreading and the length of IdU and CldU tracks was measured by immunodetection, as described previously^37,38^. This analysis showed that forks progressed at the same rate in all three cell lines (**Fig. 3b** and **Supplementary** Fig. 3a), indicating that condensin I and II are dispensable for normal fork progression in the absence of replication stress, as is the case in budding yeast.

To investigate whether condensin is required for the growth of human cells under conditions of replication stress, we focused on condensin II, which is the form present in the nucleus in interphase. The proliferation of control and CAPG2-depleted HeLa S3 cells was monitored after 3 days in the presence of various concentrations of HU (0, 100 and 300 µM). CAPG2-depleted cells had a dose-dependent growth defect in the presence of HU, ranging from 8% in the absence of HU to 58% at 300 µM HU when compared with control cells (**Supplementary** Fig. 3b). Similarly, condensin II was required for survival after exposure of the cells to 1 or 2 mM HU: CAPG2-depleted cells made ∼50% fewer colonies than control cells (**Supplementary** Fig. 3c). CAPG2-depleted cells were able to activate the DNA replication checkpoint, however, as shown by phosphorylation of the checkpoint protein CHK1 on Ser345 (**Fig. 3c**). Together, these data indicate that condensin II is required in a checkpoint-independent manner for human cells to tolerate replication stress.

To determine whether condensin promotes fork restart in human cells, as in budding yeast, we used DNA fiber spreading to quantify the percentage of restarted forks, stalled forks and newly activated origins after transient exposure to HU. HeLa S3 cells were transfected for 48 h with siRNAs against CAPG2 or with control siRNAs (siCtrl). The cells were labelled for 30 min with IdU and grown for 180 min in the presence of 4 mM HU. Then, HU was removed and the cells were labelled for a further 30 min with CldU and subjected to DNA fiber spreading (**Fig. 3d**). Strikingly, the frequency of stalled replication forks increased by 76% in CAPG2-depleted cells, and there was a concomitant 27% reduction in fork restart, which was associated with an 82% increase in the firing of new origins (**Fig. 3d**). Thus, we conclude that condensin II promotes fork restart in HeLa cells.

Our data indicate that condensin II is dispensable for normal fork progression but is important for the processing of stalled forks, as is the case in budding yeast. It has been reported that human cells respond to various sources of replication stress by ATP-dependent reversal of stressed forks to slow down replication and promote the resection of nascent DNA^39^. To determine whether condensin II is required for this replication slow down and fork resection, we labelled CAPG2-depleted cells and control cells first for 30 min with IdU and, subsequently, for 30 min with CldU in the presence of 50 µM HU, and analyzed the labelling by DNA fiber spreading. In control cells, the CldU/IdU ratio decreased by 23% upon HU treatment, indicating slowing of replication forks. Since this HU concentration is not sufficient to deplete dNTP pools, it was proposed that fork slowing under these conditions was due to fork reversal^3,40^. In CAPG2-depleted cells, by contrast, no slowing of replication was detected (**Fig. 3e** and **Supplementary** Fig. 3d), indicating that condensin II is indeed required for the slowing of DNA replication.

To determine whether condensin II is also required for fork resection, we labelled cells depleted of CAPG or CAPG2, and control cells with consecutive pulses of IdU and CldU and grew them for a further 120 min in the presence or absence of 4 mM HU. DNA fibers were stretched on glass slides, the length of IdU and CldU tracks was determined for each individual fork, and the ratio of CldU to IdU was calculated. This ratio was close to 1 for all cell types in the absence of HU, but was reduced by 26% in HU-treated control cells, reflecting the resection of nascent CldU tracks (**Fig. 3f** and **Supplementary** Fig. 3e). This limited resection activity depends on MRE11 and can be detected in various BRCA1- and BRCA2-proficient cell lines^37,38^; it differs from the hyper-resection that occurs in HU-treated, BRCA-deficient cells^41^. Remarkably, CAPG2-deficient cells were unable to resect nascent DNA, suggesting that condensin II is important for this process. Similar results were obtained in CAPG2-depleted U2OS cells – a human osteosarcoma cell line (**Supplementary** Fig. 3f). Depletion of the condensin I subunit CAPG, by contrast, did not affect fork resection (**Supplementary** Fig. 3e). These data indicate that condensin II, but not condensin I, is essential for the resection of nascent DNA at HU-arrested forks in human cells.

### Condensin II and SMARCAL1 promote fork restart by different mechanisms

Our findings indicate that condensin II promotes the slowing of replication in the presence of 50 µM HU and is essential for the resection of nascent DNA. Since both processes depend on fork reversal^3^, we investigated whether condensin II plays a direct role in this process by exposing control and CAPG2-depleted cells to 4 mM HU (**Supplementary** Fig. 4a and 4b). We also treated the cells with the MRE11 inhibitor mirin to prevent the nucleolytic degradation of reversed forks. Replication intermediates were isolated after crosslinking the DNA *in vivo* with psoralen, and visualized by electron microscopy, as described previously^42^. This analysis revealed that CAPG2-depleted cells had 2–3-fold fewer reversed forks, even in the presence of mirin, than had HU-treated control cells, in which reversed forks comprised approximately 23% of the replication intermediates observed (**Fig. 4a**, **4b** and **Table 1**), consistent with previous studies^43,44^. This indicates that condensin II facilitates fork reversal.

**Fig. 4.**
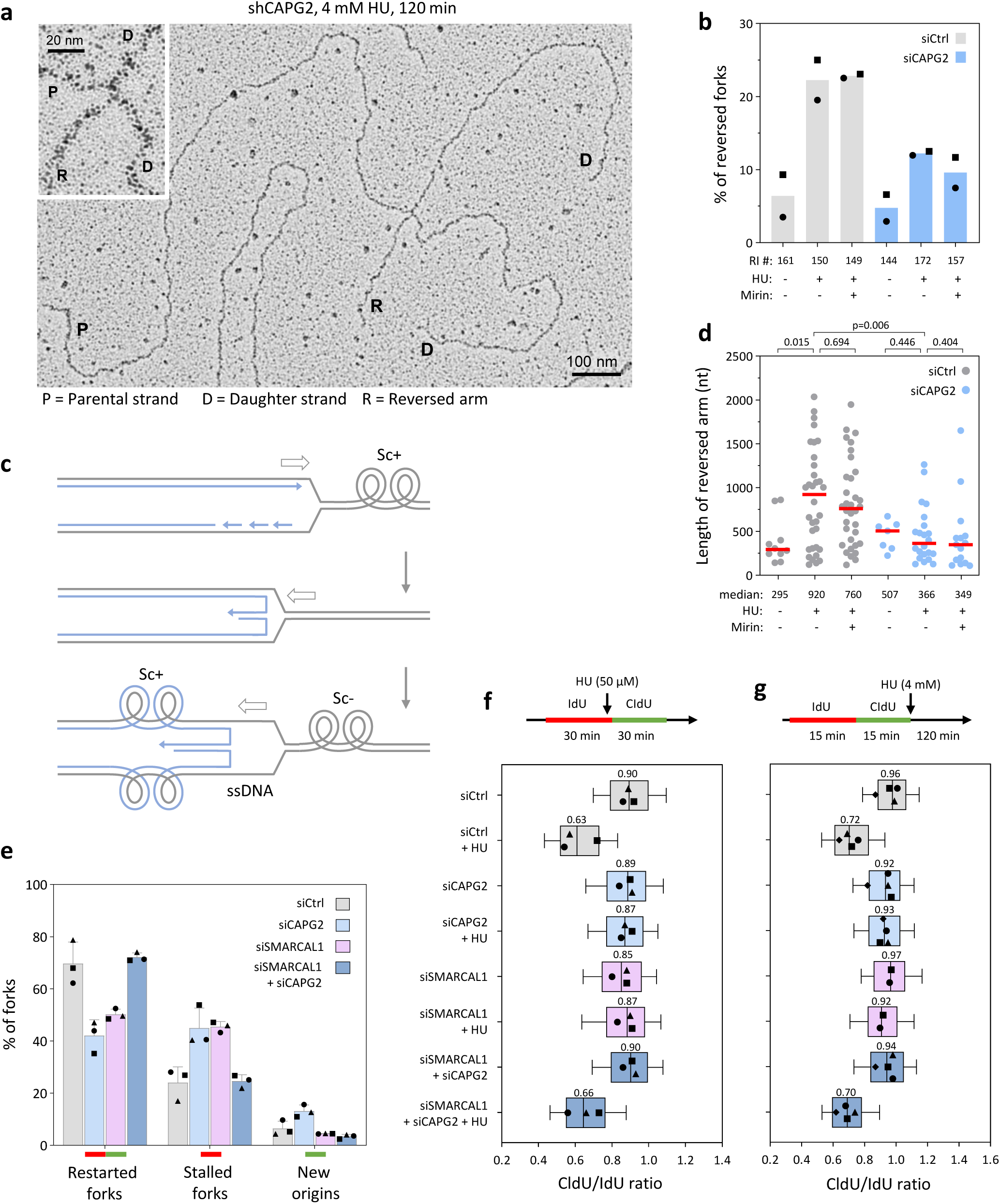
Condensin II promotes fork reversal. **a** Analysis of the frequency of reversed forks in control and condensin II-depleted cells by electron microscopy (EM). Control (shCtrl) or CAPG2-depleted (shCAPG2) HeLa-S3 cells were treated for 72 hours with 10 μM/ml doxycycline and then for 2 hours with 4 mM HU +/- 50 μM mirin before EM analysis. Electron micrographs of representative replication intermediates in shCAPG2 cells are shown. **b** The frequency of reversed forks in shCtrl and shCAPG2 cells is shown for the indicated conditions. Means and the number of analyzed forks are indicated (n=2). RI # indicates the number of analyzed replication intermediates. **c** Mechanism of fork reversal. See main text for details. **d** Reversed arms are shorter in the absence of condensin II. The length of reversed arms (in nt) was plotted for the indicated samples. Median values are indicated (n=2). RI # indicates the number of analyzed replication intermediates. P-values: Mann-Whitney rank sum test. **e** CAPG2 depletion restores fork restart in SMARCAL1-deficient cells. HeLa-S3 cells were transfected with siCtrl, siSMARCAL1, siCAPG2 or co-transfected with siSMARCAL1 and siCAPG2 for 48 hours. Cells were treated with 4 mM HU for 3 hours after a 30 minutes IdU pulse and fork restart was monitored 30 min after HU removal and CldU addition, as indicated in Fig. 3d (n=3). **f** Defective fork slowdown in SMARCAL1-deficient cells is rescued by CAPG2 depletion. U2OS cells were transfected with siCtrl, siSMARCAL1, siCAPG2 or co-transfected with siSMARCAL1 and siCAPG2 for 48 hours. Cells were labeled for 30 minutes with IdU and 30 minutes with CldU in the presence of 50 μM HU and processed for DNA fiber spreading. The ratio of CldU to IdU track length is shown for 3 independent experiments. Box and whiskers correspond to 25th–75th and 10th–90th percentiles. Median ratio is indicated. **g** CAPG2 depletion restores fork resection in SMARCAL1-depleted cells. HeLa-S3 cells were transfected with siCtrl, siSMARCAL1, siCAPG2 or co-transfected with siSMARCAL1 and siCAPG2 for 48 hours. Cells were sequentially labeled for 15 minutes with IdU and CldU, and were either collected immediately or treated for 2 hours with 4 mM HU before DNA fiber analysis. The ratio of CldU to IdU track length was plotted for four independent experiments. Box and whiskers indicate median, 25th–75th and 10th–90th percentiles.

**Table 1:** Individual EM experiments.

|  | siCtrl | siCtrl | siCtrl | siCAPG2 | siCAPG2 | siCAPG2 |
| --- | --- | --- | --- | --- | --- | --- |
|  | NT | HU | HU+Mirin | NT | HU | HU+Mirin |
| E1 | 3<br>(86) | 20<br>(82) | 23<br>(71) | 3<br>(68) | 12<br>(92) | 8<br>(80) |
| E2 | 9<br>(75) | 25<br>(68) | 23<br>(78) | 7<br>(76) | 13<br>(80) | 12<br>(77) |

\*Percentage of reversed forks – top number Number of molecules analyzed – bottom number

Replication fork reversal has been described as a two-step process^7,45^. First, a helicase, SMARCAL1, together with RAD51 and other proteins initiate limited replication fork reversal, generating positive superhelical strain in the newly replicated sister chromatids and negative superhelical strain in parental DNA. Then, TOP2A promotes extensive fork reversal by resolving the positive superhelical strain (**Fig. 4c**). In CAPG2-depleted cells, we observed not only that reversed forks were fewer, but also that the reversed arm length was two-fold shorter than in control cells (**Fig. 4d**). This is reminiscent of the phenotype of TOP2A-depleted cells^7^ and consistent with the notion that condensin II promotes the extension of fork reversal.

To characterize the mechanism by which condensin II and SMARCAL1 process stalled forks, we monitored fork arrest and restart in cells lacking CAPG2, SMARCAL1, or both proteins, by consecutive IdU and CldU labelling, as above. We found that in cells lacking either of these proteins there was an 87–89% increase in the percentage of stalled forks and a 28–40% decrease in the percentage of restarted forks (**Fig. 4e**), consistent with a role for both proteins in fork restart. When both proteins were depleted, however, the percentage of stalled forks and restarted forks was similar to those in control cells. This suggests that fork restart depends on another mechanism when SMARCAL1 and condensin II are absent.

To investigate further the roles of SMARCAL1 and condensin II in fork reversal, we used DNA fiber spreading to assay fork slowdown and fork resection in cells depleted of CAPG2, SMARCAL1, or both (**Fig. 4f, 4g** and **Supplementary** Fig. 4c–f). Like CAPG2-depleted cells, SMARCAL1-depleted cells were unable to slow replication forks (**Fig. 4f** and **Supplementary 4c**) and to resect nascent DNA (**Fig. 4g** and **Supplementary 4e**), which is consistent with a role for both proteins in fork reversal. Remarkably, depletion of CAPG2 restored fork slowdown and resection in HU-treated SMARCAL1-depleted cells (**Fig. 4f, 4g** and **Supplementary** Fig. 4c**– f**), suggesting that in the absence of condensin II, SMARCAL1 is dispensable for fork reversal.

Two RAD51-dependent fork reversal pathways have been described, which are either driven by the F-box DNA helicase FBH1 or by the combined action of SMARCAL1 and other DNA translocases^46^. To determine whether condensin II interacts functionally with FBH1, we measured fork resection by DNA fiber spreading in cells lacking FBH1, CAPG2 or both proteins. As for SMARCAL1-depleted cells, resection was abolished in the absence of FBH1 but was restored upon co-depletion of CAPG2 (**Supplementary** Fig. 4g). Moreover, the depletion of CAPG2 restored resection in cells lacking both SMARCAL1 and FBH1 (**Supplementary** Fig. 4g). Taken together, these data suggest that although the condensin II complex promotes the extension of fork reversal together with TOP2A, it may also act as a physical barrier fork reversal in the absence of SMARCAL1.

### Condensin II acts with TOP1 and TOP2A to promote fork slowing and resection

Since the positive torsional stress that accumulates ahead of the fork during DNA replication and drives fork reversal is resolved by TOP2A^6,7^, we investigated whether condensin II collaborates with TOP2A in fork reversal by assaying fork slowdown and fork resection as indicators of fork reversal in cells depleted of CAPG2, TOP2A or both proteins. TOP2A-depleted cells were defective for both fork slowing and resection (**Fig. 5a, 5b** and **Supplementary** Fig. 5a–d), consistent with its known role in fork reversal^7^ and reminiscent of the phenotype of condensin II-depleted cells. By contrast, TOP2B, which is involved in the formation of chromatin loops and TADs^8^, was dispensable for the resection of nascent DNA (**Supplementary** Fig. 5e and 5f). Cells lacking both TOP2A and CAPG2 were defective for fork slowing and resection, suggesting that these factors may act in the same pathway to promote fork reversal.

**Fig. 5.**
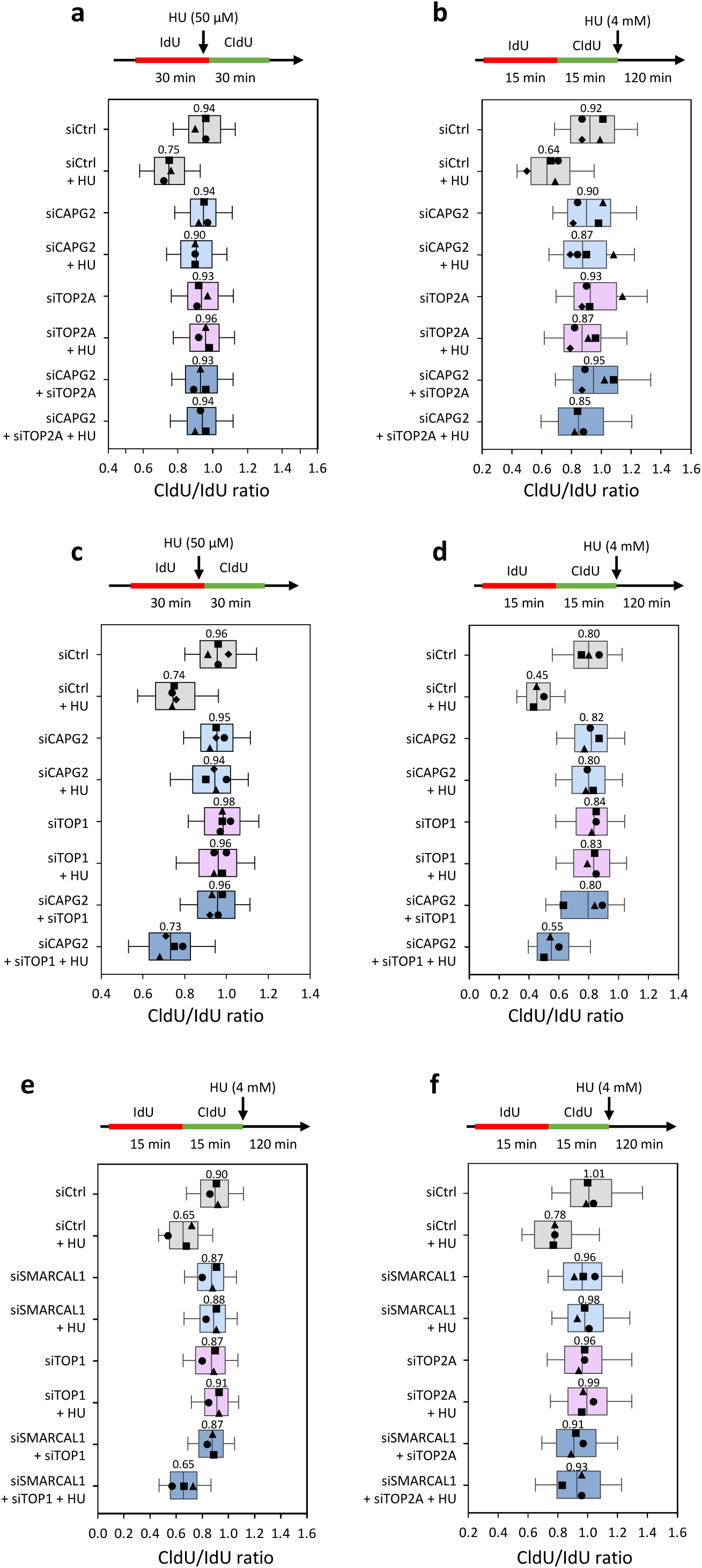
TOP1 and TOP2A play different roles in fork slowing and resection. **a** Condensin II acts with TOP2A to promote fork slowing. U2OS cells were transfected for 48 hours with siCtrl, siCAPG2, siTOP2A, or co-transfected with siTOP2A and siCAPG2. Cells were labeled for 30 minutes with IdU and 30 minutes with CldU in the presence of 50 μM HU and processed for DNA fiber spreading. The ratio of CldU to IdU track length is shown for three independent experiments. Box and whiskers correspond to median, 25th–75th and 10th–90th percentiles. **b** Condensin II acts with TOP2A to promote fork resection. HeLa-S3 cells were transfected with siCtrl, siCAPG2, siTOP2A, or co-transfected with siTOP2A and siCAPG2 for 48 hours. Cells were sequentially labeled for 15 minutes with IdU and CldU, and were either collected immediately or treated for 2 hours with 4 mM HU before DNA fiber analysis. The ratio of CldU to IdU track length was plotted for four independent experiments. Box and whiskers indicate median, 25th–75th and 10th–90th percentiles. **c** TOP1 depletion restores fork slowing in CAPG2-deficient cells. U2OS cells were transfected with siCtrl, siCAPG2, siTOP1, or co-transfected with siTOP1 and siCAPG2 for 48 hours. Cells were labeled and analyzed by DNA fiber spreading as indicated in panel a (n=4). **d** TOP1 depletion restores fork resection in CAPG2-deficient cells. HeLa-S3 cells were transfected with siCtrl, siCAPG2, siTOP1, or co-transfected with siTOP1 and siCAPG2 for 48 hours. Cells were labeled and analyzed by DNA fiber spreading as indicated in panel b (n=3). **e** TOP1 depletion restores fork resection in SMARCAL1-deficient cells. HeLa-S3 cells were transfected with siCtrl, siSMARCAL1, siTOP1, or co-transfected with siSMARCAL1 and siTOP1 for 48 h. Cells were labeled and analyzed by DNA fiber spreading as indicated in panel b (n=3). **f** Depletion of TOP2A does not restore fork resection in SMARCAL1-deficient cells. HeLa-S3 cells were transfected with siCtrl, siSMARCAL1, siTOP2A, or co-transfected with siSMARCAL1 and siTOP2A for 48 h. Cells were labeled and analyzed by DNA fiber spreading as indicated in panel b (n=3).

Since condensin activity depends on TOP1 *in vitro*^47,48^, this prompted us to investigate the role of TOP1 in fork slowing and resection. We depleted cells of CAPG2, TOP1 or both proteins and used the DNA fiber spreading assay to monitor HU-induced fork slowing and resection. These analyses showed that both processes were equally defective in the absence of TOP1 or CAPG2. Depletion of both proteins, however, restored fork slowing and resection to control levels (**Fig. 5c, 5d** and **Supplementary** Fig. 5g-j). This phenotype differs from that of cells lacking both TOP2A and CAPG2, indicating that TOP1 and TOP2A execute different functions at stalled forks.

Next, we investigated the functional interactions between SMARCAL1 and topoisomerases in fork slowing and resection. As in cells lacking CAPG2, SMARCAL1 was dispensable for fork slowing and nascent DNA resection in HU-treated TOP1-deficient cells (**Fig. 5e-f** and **Supplementary 5k** and **5i**). SMARCAL1 was required, however, to promote fork resection in the absence of TOP2A (**Fig. 5f** and **Supplementary 5m**). Thus, SMARCAL1 is dispensable for fork reversal and resection in the absence of either condensin II or TOP1, but not in the absence of TOP2A. This suggests that condensin II and TOP1 act at reversed forks to convert positively supercoiled DNA into plectonemes that are subsequently relaxed by TOP2A to promote the extension of the regressed arm.

### Condensin II and TOP1 act together to prevent accumulation of ssDNA at stalled forks

Our data show that TOP1 depletion restores fork slowing and resection in CAPG2-deficient cells (**Supplementary** Fig. 5g-j), suggesting that TOP1 has additional roles at stalled forks. Fork reversal generates not only positive supercoiling of replicated sister chromatids, but also negative supercoiling and unwinding of parental DNA. Given that TOP1 relaxes negative supercoiling and that condensin can also reanneals ssDNA^49–51^, we were prompted to investigate the role of condensin and topoisomerases in the re-annealing of parental DNA strands during fork reversal.

To this end, we first assayed the abundance of RPA-coated ssDNA at HU-arrested forks in yeast cells by ChIP-qPCR. Wild type, *SMC4*-AID, *top1Δ* and *top1Δ SMC4-AID* cells were released synchronously from G_1_ into S phase for 60 min in the presence of 200 mM HU and RPA-coated ssDNA was immunoprecipitated and analyzed by qPCR, as described previously^27^. More RPA was found at forks progressing from the early origins *ARS306* and *ARS607* in *SMC4*-AID cells than was found in control cells (**Fig. 6a**). A similar increase in the amount of RPA at *ARS607* was also found in *YCG1*-AID cells depleted of the condensin subunit Ycg1 (**Supplementary** Fig. 6a). Remarkably, yet more RPA was found at *ARS306* and *ARS607* in *top1Δ* and *top1Δ SMC4-AID* cells (**Fig. 6a**), and at forks progressing from *ARS305* (**Supplementary** Fig. 6b). These data support the notion that condensin and Top1 act together to prevent the accumulation of RPA-coated ssDNA at HU-arrested forks in budding yeast.

**Fig. 6.**
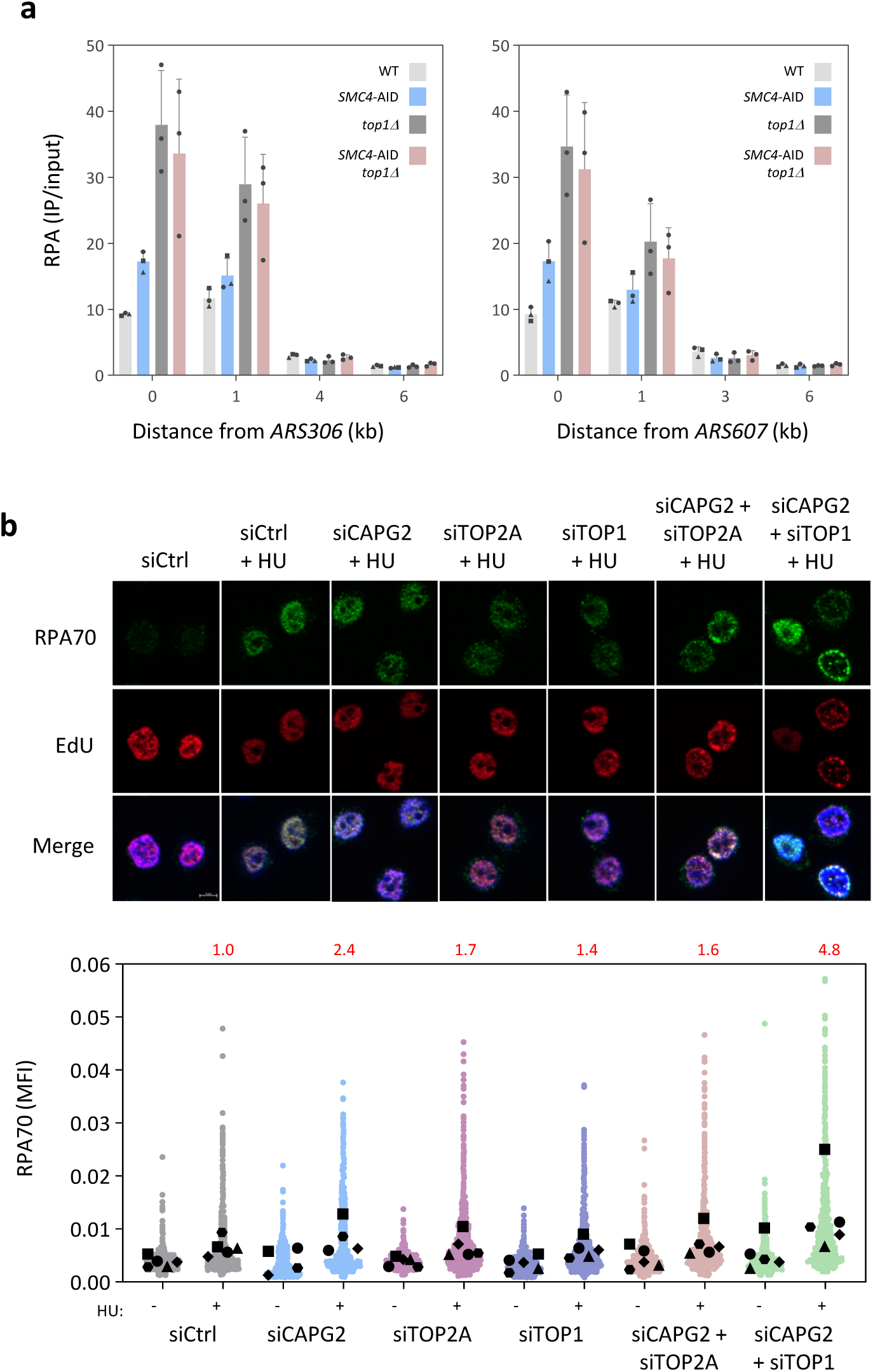
Condensin II and TOP1 act together to prevent accumulation of ssDNA at stalled forks. **a** RPA-coated ssDNA accumulates at HU-arrested forks in yeast cells lacking condensin and Top1. WT, *SMC4*-AID, *top1D* and *top1D SMC4*-AID cells were arrested in G_1_ with a-factor and SMC4 was depleted by addition of auxin for 60 minutes. Then, cells were released into S phase in the presence of 200 mM HU for another 60 minutes. RPA enrichment at indicated distances from *ARS306* and *ARS607* was determined by ChIP-qPCR after normalization to unreplicated regions. Mean and SD correspond to three independent experiments. **b** Levels of chromatin-bound RPA increase in the absence of both CAPG2 and TOP1. HeLa-S3 cells were transfected for 48 hours with siCtrl, siCAPG2, siTOP2A, siTOP1 or co-transfected as indicated. Cells were labeled with EdU and were treated with 4mM of HU for 2 hours. RPA70 (green) and EdU-labelled replication sites (red) were immunodetected. The mean fluorescence intensity (MFI) of RPA70 was quantified in 5 independent experiments using CellProfiler. Changes in median intensity relative to control cells are indicated in red.

To determine whether this is also the case in human cells, we assayed the accumulation of RPA foci by immunofluorescence microscopy of human cells depleted of either CAPG2, TOP1, TOP2A or various combinations of these enzymes. This analysis revealed a 2.4-fold increase in the fluorescence intensity of RPA foci upon HU addition in CAPG2-depleted cells when compared with control cells (**Fig. 6b**). This increased further to 4.8-fold in cells depleted of both CAPG2 and TOP1, but not in cells depleted of CAPG2 and TOP2A (**Fig. 6b**). We conclude, therefore, that condensin acts with TOP1 to prevent the formation of excess RPA-coated ssDNA at HU-arrested forks.

## DISCUSSION

In this study, we investigated the role of condensin in S phase and, specifically, whether condensin and topoisomerases cooperate to relieve positive DNA supercoiling at replication stress sites. We show that condensin is recruited to stalled forks in yeast by the MRX complex, where it cooperates with the topoisomerases Top1 and Top2 to promote fork restart. Also, in human cells, we find specifically condensin II is essential for resection of nascent DNA at stalled forks, allowing them to restart replication. Condensin II and SMARCAL1 execute different functions to promote fork restart in human cells. Condensin II also cooperates with topoisomerases at stalled forks: with TOP1 to prevent accumulation of ssDNA and with TOP1 and TOP2A to promote fork reversal. Our findings are consistent with a model in which condensin II and TOP1 act at reversed forks to convert positively supercoiled DNA into structures that are subsequently relaxed by TOP2A allowing the fork to resume replication.

### Condensin is recruited to replication forks in yeast and human cells

Condensin is reported to bind multiple loci in the DNA of interphase *S. cerevisiae*, including replication origins and transcription sites^12^. Here, we found condensin enriched specifically around active early replicating origins, not at repressed late origins in HU-treated *S. cerevisiae* cells, indicating that it binds origins after initiation. This is reminiscent of condensin recruitment to centromeric regions of *S. cerevisiae* chromosomes, which also depends on DNA replication^14^. In HU-treated *S. cerevisiae* cells, replication forks progress very slowly (0.1 kb per minute)^24^, allowing us to track the position of condensin in time-resolved ChIP-seq experiments. We found that condensin moved away from origins with the same kinetics as DNA replication, suggesting that it follows stressed replication forks. In human cells, our iPOND analysis showed that condensin associates with newly replicated DNA in the absence of HU^52^, which is consistent with earlier results^53^ and suggests that active forks can also recruit condensin.

Our findings raise the question of how condensin is recruited to replication sites. Condensin is reported to be recruited to transcriptionally active chromatin regions by transcription factors and nucleosome displacement^54–57^, but the mechanism involved remains unclear. In HU-treated yeast cells, we found that MRX-dependent remodeling of nascent chromatin is an important determinant of condensin recruitment to replication sites, as described earlier for cohesin^27^. Our data do not rule out the possibility that condensin could also be loaded elsewhere in the genome and slide along the DNA until it encounters barriers such as tightly bound protein complexes^58–60^, closed topological domains^5,6^ or entangled DNA structures^33,61^. Whether condensin complexes recruited to replication pause sites in S phase contribute to chromosome condensation in G_2_/M phase remains an important question to address.

### A role for condensin in fork repair

We show that loss of condensin from yeast and human cells reduced cell growth and survival in response to replication stress. This is consistent with earlier studies, showing that condensin mutations increase sensitivity to DNA damage in several species^17–22,28^. Together, these findings suggest either that condensin plays a direct role in DNA repair in interphase or that DNA damage indirectly increases chromosome segregation defects, which are frequent in condensin-deficient cells. To determine whether condensin plays a direct role in DNA replication and repair, we depleted condensin in G_1_-arrested *S. cerevisiae* cells and used DNA combing to monitor fork progression, arrest and restart in the subsequent S phase. This analysis revealed that condensin was required for the timely recovery of stressed forks after HU- or MMS treatment, independently of its classical function in chromosome segregation. This phenotype was also independent of the activation of the S-phase checkpoint, unlike in fission yeast^28^. Condensin II depletion in human cells also impaired fork restart without affecting checkpoint activation, suggesting that condensin plays a direct role in fork repair.

### Condensin acts with TOP2A to promote fork reversal in human cells

We used three independent experimental approaches in this study to show that condensin promotes fork reversal in human cells. Depletion of the condensin II subunit CAPG2 reduced both fork resection and slowdown in HU-treated cells, which are two important consequences of fork reversal^3^. Also electron microscopy of replication intermediates confirmed that fork reversal is impaired in condensin II-depleted cells. Interestingly, the regressed arms were not only less abundant than in control cells but were also shorter, suggesting that condensin II is involved in the extension step of fork reversal. Indeed, fork reversal is a two-step process consisting of the initial formation of an unstable short reversed arm and its subsequent extension through a process involving TOP2A, the SUMO E3 ligase ZATT, and a SUMO-targeted DNA translocase, PICH^3,7^. TOP2A promotes the extension of reversed forks by relaxing positive DNA supercoiling that accumulates upon unwinding of sister chromatid DNA by SMARCAL1 and other translocases. The co-depletion of CAPG2 and TOP2A did not further impede fork resection and slowdown, suggesting that condensin II and TOP2A act in the same pathway to promote fork reversal.

This collaboration between condensin and TOP2A is reminiscent of their role in M phase^9^. TOP2A catalyzes the catenation and decatenation of sister chromatids in G_2_/M phase, and condensin promotes chromosome segregation by tilting the balance towards decatenation^32,33,62–64^. Inactivation of condensin in metaphase results in *de novo* formation of sister chromatid intertwines^65,66^, suggesting that condensin promotes the unlinking of sister chromatids by TOP2A whereas their close proximity favors interlinking. The mechanism involved, however, remains a subject of debate. Condensin might drive TOP2 decatenation activity by inducing the formation of positive supercoiling^31,65^ or by constricting intra- and inter-molecular DNA contacts without altering DNA supercoiling^33,67^. It might also bind positive superhelical plectonemes and extrude them into a stable supercoiled loop to prevent the formation of harmful DNA knots and sister chromatid intertwines^68^. We speculate that condensin II might also sequester positive superhelical DNA into large plectonemes during fork reversal, which would both favor their relaxation by TOP2A and prevent the formation of harmful topological structures (**Fig. 7**). This activity would differ from its mitotic role when condensin drives the formation of positive supercoils in anaphase^14,47^, whereas in S phase positive supercoiling is generated by fork reversal in conditions of replication stress.

**Fig. 7.**
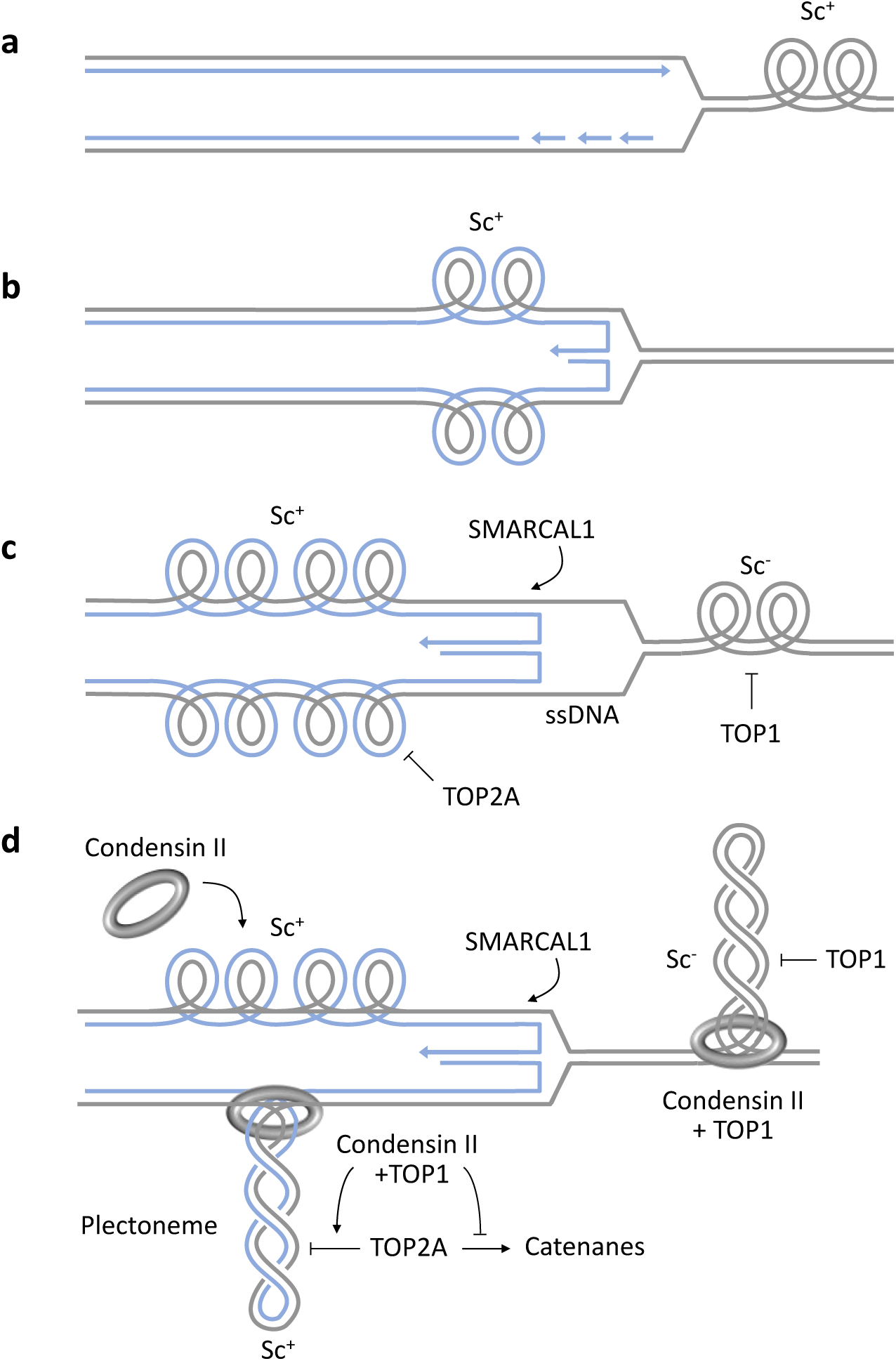
Model of the roles of condensin and topoisomerases in fork reversal. **a** Elongation of DNA replication generates positive supercoils (Sc+) in front of the forks that are normally relaxed by TOP1 and precatenanes behind the fork that are resolved by TOP2. **b** Upon fork arrest, Sc+ can be transferred behind the fork via a process called fork reversal, in which nascent DNA strands reanneal to form a regressed arm. In a topologically closed context, Sc+ represents a topological barrier that limits the extent of fork reversal. **c** Further fork reversal is mediated by DNA translocases such as SMARCAL1, which induce additional Sc+ along newly replicated sister chromatids. Fork reversal also increases Sc-levels and ssDNA on parental DNA. Fork reversal promotes fork slowing and generates a one-ended DNA arm that is susceptible to nucleolytic degradation. **d** Condensin II binds Sc+ and drives the formation of positively supercoiled plectonemes together with TOP1. These plectonemic structures define topologically closed domains that promote the relaxation activity of TOP2 and prevent the formation of catenanes. In the absence of condensin II or TOP1, these topological barriers do not form and TOP2 is sufficient to promote fork reversal in a SMARCAL1-independent manner, at the expense of increased fork restart defects. Finally, our data indicate that condensin acts on parental DNA to prevent the accumulation of ssDNA and promote the relaxation of Sc-by TOP1.

### Cooperation between condensin II and SMARCAL1 during fork reversal

It has been proposed that TOP2A, ZATT, and PICH promote fork reversal by relaxing positive supercoiling generated by SMARCAL1 and other translocases^7^. Our data are consistent with this model as the depletion of either TOP2A, SMARCAL1 or both enzymes equally prevented fork slowdown and resection. We also found that the depletion of both condensin II and SMARCAL1 restored fork slowdown and resection in HU-arrested cells, suggesting that SMARCAL1 is dispensable for fork reversal in the absence of condensin. This finding is surprising as SMARCAL1 is generally believed to be essential for fork reversal. A possible interpretation of these data is that the plectonemes generated by condensin during fork reversal act as a physical barrier for the extension of the reversed arm. In the absence of this barrier, the annealing activity of SMARCAL1 might be dispensable for fork reversal or may be mediated by other factors, such as RAD51^3,69^.

### TOP1 and TOP2A promote fork reversal by different mechanisms

TOP1 is also a key enzyme involved in the resolution of negative and positive superhelical tension^8^. We found that, like TOP2A and CAPG2, TOP1 is essential for fork slow down and reversal at stalled forks. Unlike TOP2A, however, depletion of TOP1 rescued fork defects in SMARCAL1-depleted cells. Since *in vitro* TOP1 relieves the torsional stress generated when condensin induces positive supercoiling of DNA by trapping supercoils^34,47,70^, it might be essential *in vivo* for the function of condensin at reversed forks. Our finding that depletion of both TOP1 and CAPG2 rescued the replication defects observed when either of these proteins was depleted, suggests that TOP1 has another role in fork reversal in addition to relaxation of positive superhelical tension. The negative superhelical tension generated at reversed forks by reannealing of parental DNA strands is also a good substrate for TOP1 (**Fig. 7**). In TOP1-depleted yeast and human cells, we observed accumulation of RPA-coated ssDNA in response to HU, which would be consistent with a role for TOP1 in promoting the reannealing of parental DNA. Interestingly, increased ssDNA levels were also detected in cells lacking condensin and TOP1, consistent with a role for these factors in ssDNA reannealing (**Fig. 7**), as reported for replication of centromeric DNA in *Xenopus* egg extracts^50^.

Taken together, our findings paint a complex picture in which condensin works in concert with DNA topoisomerases to relieve topological constraints generated during fork reversal while preventing the formation of harmful knots and sister chromatid intertwines by TOP2A. They raise important questions about the topological structures generated by condensin II at reversed forks, and about the coordinated activity of condensin and TOP1 ahead of and behind reversed forks. Since other SMC complexes – cohesin, MRX and its human homolog MRN, and SMC5/6 – are also recruited to stalled forks under replication stress conditions^27,71,72^, it will be important to investigate the coordination between condensin and these other SMCs during fork reversal. In particular, our data indicate that MRX is required to recruit condensin to HU-arrested forks in budding yeast, as is the case for cohesin^27^. Since cohesin and condensin cooperate in M phase to ensure proper chromosome segregation, this might also be the case at reversed forks. Although condensin promotes fork restart in yeast and human cells, the mechanisms involved may differ as other recombinational repair mechanisms such as template switching are also very active in budding yeast^73,74^. Finally, an important question to be addressed is whether the condensin-mediated processes observed in HU-treated cells also operate during normal S phase, in the absence of replication stress, and, if so, to what extent they contribute to the loading of condensin II in interphase to drive chromosome condensation and segregation in M phase.

## MATERIALS AND METHODS

### Standard yeast genetics

All strains used were listed in the **Table 2**. All strains used were haploid and derived from W303. For the cell sensitivity to genotoxic drugs, cells were adjusted to 1x10^7^ cells/mL and serial dilution (1:10) were spotted on plates with HU (Sigma, H8627) or MMS (Sigma, 129925).

**Table 2:** Yeast strains used in this study.

| Name | Genotype | Figure |
| --- | --- | --- |
| PP870 | MATa, ade2-1, trp1-1, can1-100, leu2-3,112, his3-11,15, ura3, GAL, psi+, RAD5, flo11 A2758G | 1A |
| PP872 | MATa, ade2-1, trp1-1, can1-100, leu2-3,112, his3-11,15, ura3, GAL, psi+, RAD5, URA3::GPD-TK7 | 1B, 1C, 1D, S1B, S1C |
| PP1919 | MATa, ade2-1, trp1-1, can1-100, leu2-3,112, his3-11,15, ura3, GAL, psi+, RAD5, URA3::GPD-TK7 Brn1-PK9::TRP | 1B, 1C, 1D, 1E, S1B, S1C |
| PP1941 | MATa, ade2-1, trp1-1, can1-100, leu2-3,112, his3-11,15, ura3, GAL, psi+, RAD5, URA3::GPD-TK7 Brn1-PK9::TRP, rad50::KAN | S1B, S1C |
| PP2041 | MATa, ade2-1, trp1-1, can1-100, leu2-3,112, his3-11,15, ura3, GAL, psi+, RAD5, smc2-8::KAN | 1A |
| PP2103 | MATa, ade2-1, trp1-1, can1-100, leu2-3,112, his3-11,15, ura3, GAL, psi+, RAD5, Brn1-PK9::TRP | 1F |
| PP2405 | MATa, ade2-1, trp1-1, can1-100, leu2-3,112, his3-11,15, ura3, GAL, psi+ , RAD5, TK-hENT1::LEU2, ADH1pr-O.s.Tir1-myc9::TRP1 | 6A, S6B |
| PP2408 | MATa, ade2-1, trp1-1, can1-100, leu2-3,112, his3-11,15, ura3, GAL, psi+ , RAD5, TK-hENT1::LEU2, ADH1pr-O.s.Tir1-myc9::TRP1, SMC4-3PK-1xminiAID-KAN | 6A, S6B |
| PP2668 | MATa, ade2-1, trp1-1, can1-100, leu2-3,112, his3-11,15, ura3, GAL, psi+ , RAD5, , ADH1pr-O.s.Tir1-myc9::TRP1, SMC4-3PK-1xminiAID-KAN, top1::NAT | 6A |
| PP2670 | MATa, ade2-1, trp1-1, can1-100, leu2-3,112, his3-11,15, ura3, GAL, psi+ , RAD5, , ADH1pr-O.s.Tir1-myc9::TRP1, top1::NAT | 6A |
| PP3136 | MATa, ade2-1, trp1-1, can1-100, leu2-3,112, his3-11,15, ura3, GAL, psi+, RAD5, Brn1-PK9::TRP, gcn5::HIS | 1E |
| 3601 | MATa, ade2-1, trp1-1, can1-100, leu2-3,112, his3-11,15, ura3, GAL, psi+, RAD5, Brn1-PK9::TRP, chd1::TRP | 1F |
| PP4947 | MATa,ade2-1, trp1-1, can1-100, leu2-3,112, his3-11,15, ura3, GAL, psi+, RAD5+, ade2-1::OsTIR1-9myc:ADE2 ura3-1::ADH1-OsTIR1-2-9Myc (URA3) SMC4-3Pk-miniAID:kanR, AUR1::yTK-yENT1::AUR1C | S1A, S2A, S2B, 2C, 2D, S2F, S2G, S2H, S2I, S2J |
| PP4948 | MATa,ade2-1, trp1-1, can1-100, leu2-3,112, his3-11,15, ura3, GAL, psi+, RAD5+, ade2-1::OsTIR1-9myc:ADE2 ura3-1::ADH1-OsTIR1-2-9Myc (URA3) SMC4-3Pk-miniAID:kanR, trp1-1::PSMC4-SMC4WT-3HA:TRP1, AUR1::yTK-yENT1::AUR1C | S1A, S2A, 2B, S2F |
| PP4949 | MATa, leu2-3,112 his3-11,15 ade2-1 trp1-1 can1-100 RAD5+ ura3-1::pADH1-OsTIR1-URA3, ade2-1::OsTIR1-9myc:ADE2, TOP2-AID*-MYC-HPH, AUR1::yTK-yENT1::AUR1C | 2D, S2H, S2J |
| PP4950 | MATa, leu2-3,112 his3-11,15 ade2-1 trp1-1 can1-100 rad5-535<br>ura3-1::pADH1-OsTIR1-URA3, ade2-1::OsTIR1-9myc:ADE2, ,<br>AUR1::yTK-yENT1::AUR1C | 2B, 2D, S2F |
| PP4952 | MATa, leu2-3,112 his3-11,15 ade2-1 trp1-1 can1-100 RAD5+<br>ura3-1::pADH1-OsTIR1-URA3, ade2-1::OsTIR1-9myc:ADE2, TOP2-<br>AID*-MYC-HPH, SMC4-3Pk-miniAID:kanR, , AUR1::yTK-<br>yENT1::AUR1C | 2D, S2H, S2J |
| PP5214 | MATa, leu2-3,112 his3-11,15 ade2-1 trp1-1 can1-100 rad5-535<br>ura3-1::pADH1-OsTIR1-URA3, ade2-1::OsTIR1-9myc:ADE2,<br>AUR1::yTK-yENT1::AUR1C | 2C, S2G, S2I |
| PP5481 | MATa, ade2-1, trp1-1, can1-100, leu2-3,112, his3-11,15, ura3, GAL,<br>psi+, RAD5+, YCG1-3PK:: K.lactis TRP1, AUR1::yTK-yENT1::AUR1C | S2A, S2B, S2D, S2E,<br>S6A |
| PP5579 | MATa, ade2-1, trp1-1, can1-100, leu2-3,112, his3-11,15, ura3, GAL,<br>psi+, rad5+, (K.lactis URA3)pMET3-YCG1-3PK-miniAID::KANMX6,<br>ADE2::pADH1-OsTIR1-9Myc, AUR1::yTK-yENT1::AUR1C | S2A, S2B, S2D, S2E,<br>S6A |
| PP5589 | MATa,ade2-1, trp1-1, can1-100, leu2-3,112, his3-11,15, ura3, GAL,<br>psi+, RAD5+, ade2-1::OsTIR1-9myc:ADE2 ura3-1::ADH1-OsTIR1-2-<br>9Myc (URA3) SMC4-3Pk-miniAID:kanR TOP1-AID-KanMX6?<br>AUR1::yTK-yENT1::AUR1C | 2C, S2G, S2I |
| PP5599 | MATa,ade2-1, trp1-1, can1-100, leu2-3,112, his3-11,15, ura3, GAL,<br>psi+, RAD5+, ade2-1::OsTIR1-9myc:ADE2 ura3-1::ADH1-OsTIR1-2-<br>9Myc (URA3) TOP1-AID-KanMX6 AUR1::yTK-yENT1::AUR1C | 2C, S2G, S2I |

### Cell growth and synchronization

Cells were grown at 25°C in YPD or in YN-without methionine, supplemented with 2% glucose. 6x10^6^ exponentially growing cells were synchronized in G_1_ using 6 μg/mL of α-factor (Biotem, 2968) during 170min. G_1_ arrested cells were released into S phase by filtration and resuspension in fresh medium or by the addition of 75 μg/mL Pronase (Sigma, 53702). Cells were treated or not with 200mM HU or 0.05% MMS, as indicated. For degron strains, 1 mM auxin (IAA) was added to the liquid culture after 120 min of synchronization in G_1_ and every hour after release in S phase.

### Human cell culture

Human cervical adenocarcinoma HeLa S3 cells (ATCC, CCL2-2) and osteocarcinoma U2OS cells (ATCC, HTB-86) were cultured in Dulbecco’s modified Eagle’s medium (DMEM) or McCoy’s 5a medium, respectively, supplemented with 10% fetal calf serum (FCS) and 100 U/mL penicillin/streptomycin (PS). HeLa-S3 shCtrl and shCAPG2 were cultured in DMEM supplemented with 10% tetracyclin-free FCS and 100 U/mL PS. All cells were cultured at 37°C in a 5% CO2 incubator.

### Production of lentiviral vectors and transduction

HIV-1-derived lentiviral vectors were produced in HEK293T cells as previously described^75^. Cells were seeded on poly-D-lysine coated plates and transfected with packaging plasmid (psPAX2, Addgene plasmid #12260): transfer vector (pLVX-Tet-on; TRIPZ-shCtrl, TRIPZ-shCAPG2): vesicular stomatitis virus envelop plasmid (pMD2.G, plasmid #12259) at a ratio 5:3:2 by the calcium phosphate method. The culture medium was collected 48h post-transfection, filtrated using 0.45 µm filters and concentrated at 100-fold by ultracentrifugation at 89,000 g at 4 °C for 1h30. HeLa cells were transduced at a M.O.I = 10 (Multiplicity of Infection) by centrifugation at 1500 g at 30°C for 90 minutes in the presence of 5 µg/ml of polybrene.

### Cell transfection and RNA interference

HeLa-S3 and U2OS cells were transfected for 48 hours using the lipofectamine RNAiMAX reagent and siRNAs as indicated in **Table 3**. HeLa-S3 shCtrl and shCAPG2 were selected with puromycin (1 μg/mL) and shRNAs were induced with 4 μg/mL of Doxycycline (D3072) for 72 hours.

**Table 3:**
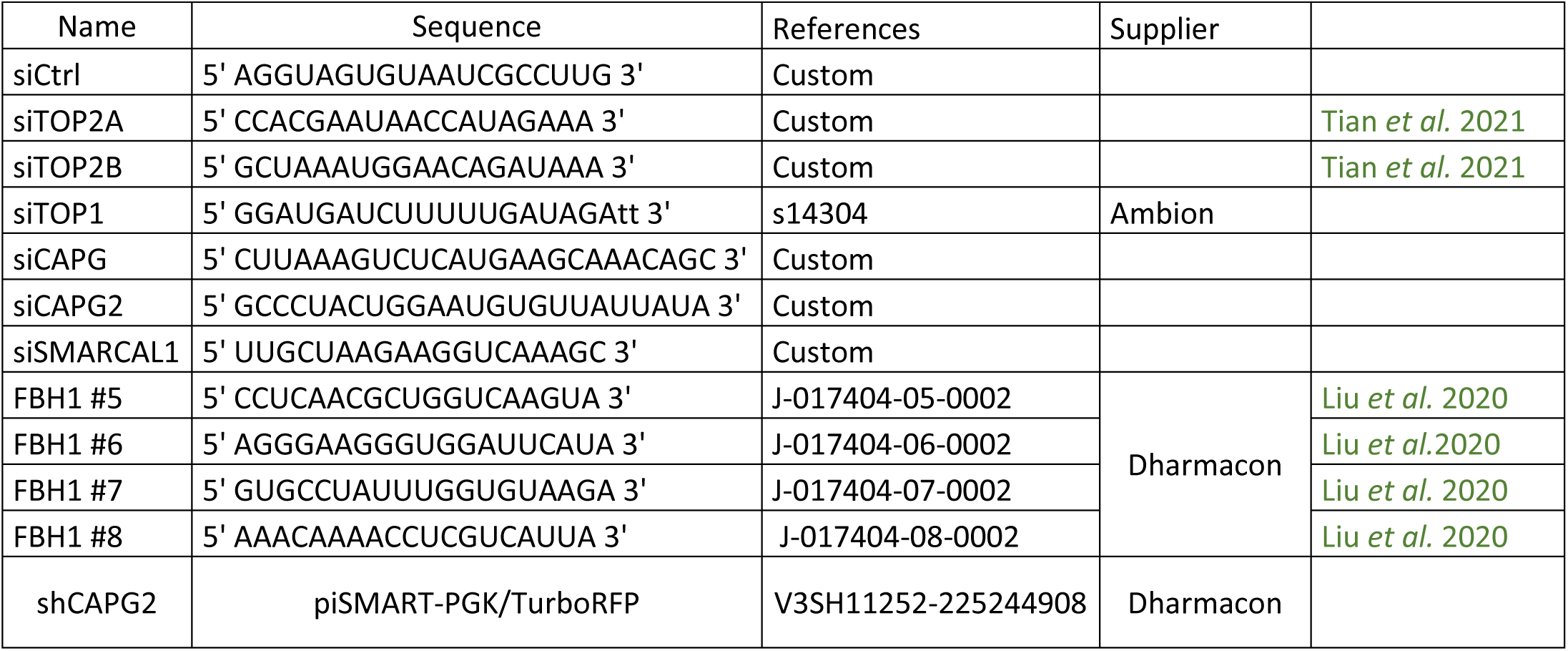
siRNA or shRNA used in this study.

**Table 4:**
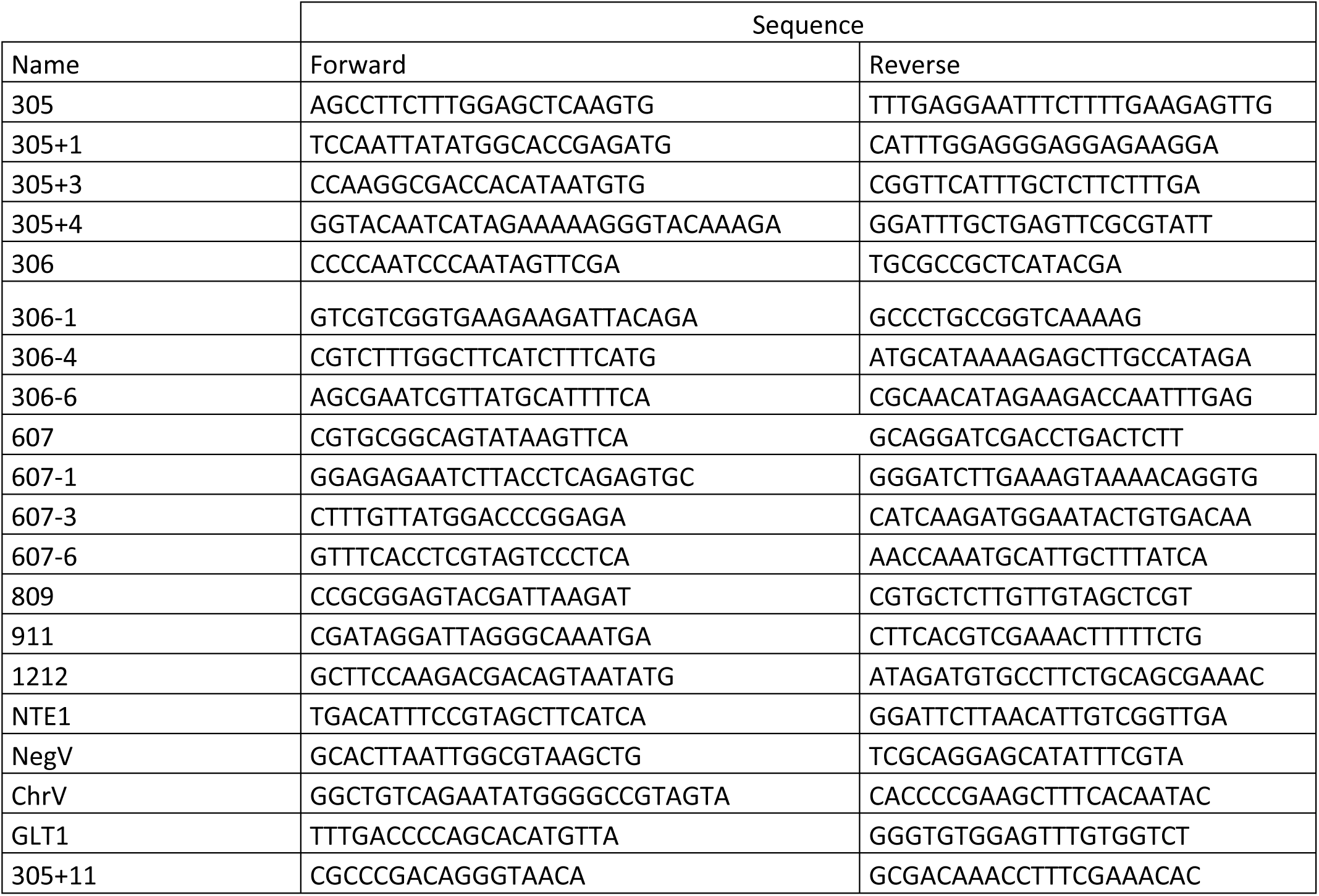
Oligonucleotides used in this study.

### Flow cytometry

Yeast cells: 500 μL of cells were fixed with 1 mL of ethanol 100%. Cells were centrifuged for 1min at 16 000 RCF, resuspended in 500 μl of 50mM Na-citrate pH=7.5 containing 10 μL of RNase A (10 mg/mL) and incubated 2 hours at 50°C. Then, 10 μL of proteinase K (20 mg/mL) were added and incubated 2 hours at 50°C. Aggregates of cells were dissociated by sonication. 30 μL of cells were incubated with 270 μL of 50 mM Na-citrate pH=7.5 containing 1X of Sytox (10 000X) for label DNA.

Human cells: Cells were labelled with 10 μM EdU for 20min. For the fixation, cells were incubated with 1% formaldehyde/PBS in the dark during 10min at room temperature and washed in 5% BSA in PBS. For the permeabilization, cells were incubated 30min in the dark at room temperature with 0.2% Triton X-100 in 1% BSA in PBS. Click reaction (PBS, 3mM CuSO4, 29 μM A488 azide dye, 10 mM sodium ascorbate) was performed during 30min at room temperature in the dark. Cells were then incubated for 30min with 0.1 mg/mL RNase and 2 μg/mL DAPI.

For both type of experiments, data were acquired on a MACSQuant and analyzed with FlowJo software.

### Chromatin Immunoprecipitation

Chromatin immunoprecipitation was performed as described in Delamarre *et al*, 2020^27,71^. 5x10^8^ cells were crosslinked for 15 min with 1% formaldehyde (Sigma F8775) at RT under agitation. Fixation was quenched by addition of 0.25 M Glycine (Sigma G8898) for 5 min under agitation. Cells were washed three times with cold TBS1X (4°C). Dry pellets were frozen and conserved at -20°C. Cell pellets were resuspended in 600 µL lysis buffer (50 mM HEPES-KOH pH7.5, 140 mM NaCl, 1 mM EDTA, 1% Triton X-100, 0.1% Na-deoxycholate) supplemented with 1 mM PMSF and anti-protease (cOmplete Tablet, Roche, 505649001) and lysed by beads-beat method (MB400 U, Yasui Kikai, Osaka). Recovered lysate (WCE, Whole Cell Extract) was sonicated with a Q500 sonicator Bioruptor (4 cycles: 30 s ON, 30 s OFF). Dynabeads were washed three times and resuspended in 1 mL of PBS, 0.5% BSA, 0.1% Tween and incubated with antibodies on a rotating wheel for two hours at 4°C. 2.5 µL of anti-RPA (Agrisera, AS07214) with 90 µL Dynabeads Prot. A (DPA). 20 µl of anti-PK (Anti-V5 tag, AbD Serotec, MCA1360G) with 90 µl DPA. 25 µl of WCE were kept for the Input sample and 25 µl were collected for western-blotting (WB). Antibodies coupled Dynabeads were washed three times with 1 mL of PBS, 0.5% BSA, 0.1% Tween and added to 500 µL of WCE overnight on a rotating wheel at 4°C. Beads were then collected on a magnetic rack. 25 µl of the supernatant were collected for WB analysis (Flow-Through sample) and beads were washed on ice with cold solutions: two times with Lysis buffer (50 mM HEPES-KOH pH7.5, 140 mM NaCl, 1 mM EDTA, 1% Triton X-100, 0.1% Na-deoxycholate), twice with Lysis buffer + 0.36 M NaCl (50 mM HEPES-KOH pH7.5, 360 mM NaCl, 1 mM EDTA, 1% Triton X-100, 0.1% Na-deoxycholate), twice with Wash buffer (10 mM Tris-HCl pH8, 0.25 M LiCl, 0.5% IGEPAL, 1 mM EDTA, 0.1% Na-deoxycholate) and once with TE (10 mM Tris-HCl pH8, 1 mM EDTA). Antibodies were uncoupled from beads with 120 µl of Elution Buffer (50 mM Tris-HCl pH8, 10 mM EDTA, 1% SDS) for 10 min at 65°C. 5 µl of eluates were collected for WB (IP sample) and 115 µl were incubated with 120 µl of TE, 0.1% SDS for de-crosslinking at 65°C overnight. 130 µl of TE containing 60 mg RNase A (Sigma, R65-13) were added to the samples and incubated for 2 hours at 37°C. Proteins were digested by addition of 20 µl of Proteinase K (Sigma, P6556) at 20 mg/ml and incubated for 2 hours at 37°C. 50 µL of 5M LiCl were added to DNA before purification by two rounds of Phenol: Chloroform: Isoamyl Alcohol 25:24:1 (Sigma, P2069) extractions and precipitation by addition of 100 mM Sodium Acetate (Sigma, S2889), 26 mg/ml of Glycogen (Roche, 10901393001) and two volumes of 100% ethanol overnight at -20°C. Samples were centrifuged for 45 min at 16.000 RCF at 4°C, washed with cold 70% ethanol and centrifuged again 15 min at 16.000 RCF at 4°C. DNA pellets were dried and resuspended in 300 µl of H20 prior to qPCR reactions or in 22 µL prior to deep-sequencing. qPCR reaction was performed with LightCycler480 (Roche). IP/Input ratio were calculated and qPCR results were normalized with five negative zones: ChrV, NegV, NTE1, 305+11, GLT1. Aspecific IP and western-blotting of the protein samples were performed for each experiment.

### Genome-wide profiling

Genomic DNA was isolated using Qiagen genomic DNA extraction kit. DNA was fragmented using sonication (200 to 500 bp). ChIP sequencing libraries were prepared using the NEBNext Ultra II DNA Library Prep Kit for Illumina (NEB). Next generation sequencing was performed on a NovaSeq 6000 (Illumina). Single-end reads of 50 bp and paired-end reads were aligned to *S. cerevisiae* genome (2011) sequence with Bowtie/2. ChIP-seq profiles expressed as RPKM were obtained as a ration of IP/input reads. Relative DNA content was determined as a ratio of normalized HU reads to G_1_ reads.

### Western botting

Yeast cells: Total protein extracts extraction was performed as described in Poli et al. 2016. Human cells: cells pellets were resuspended in 2X Laemmli buffer and treated with Benzonase (25 U/μL) for 30 min at 37°C twice.

Proteins were resolved by SDS-PAGE (Biorad) and then transferred to nitrocellulose membranes. After blocking, proteins were probed with antibodies listed in **Table 3**. Membranes were scanned with a ChemiDoc MP (Biorad).

### DNA Fiber Spreading and DNA Combing

DNA combing was performed as described^30^. For BrdU labelling detection, a rat monoclonal anti-BrdU was used. A click-it reaction kit was used for EdU labelling detection. A mouse monoclonal anti-ssDNA was used to detect DNA fibers.

DNA fiber spreading was performed as described previously^37,38^. Briefly, subconfluent cells were sequentially labeled with 10 µM 5-iodo-2’-deoxyuridine (IdU) and with 100 µM 5-chloro-2’-deoxyuridine (CldU) for the indicated times. One thousand cells were loaded onto a glass slide (StarFrost) and lysed with spreading buffer (200 mM Tris-HCl pH 7.5, 50 mM EDTA, 0.5% SDS) by gently stirring with a pipette tip. The slides were tilted slightly and the surface tension of the drops was disrupted with a pipette tip. The drops were allowed to run down the slides slowly, then air dried, fixed in methanol/acetic acid 3:1 for 10 minutes, and allowed to dry. Glass slides were processed for immunostaining with mouse anti-BrdU to detect IdU, rat anti-BrdU to detect CldU, mouse anti-ssDNA antibodies and corresponding secondary antibodies conjugated to various Alexa Fluor dyes. Antibodies are indicated in **Table 5**.

**Table 5:**
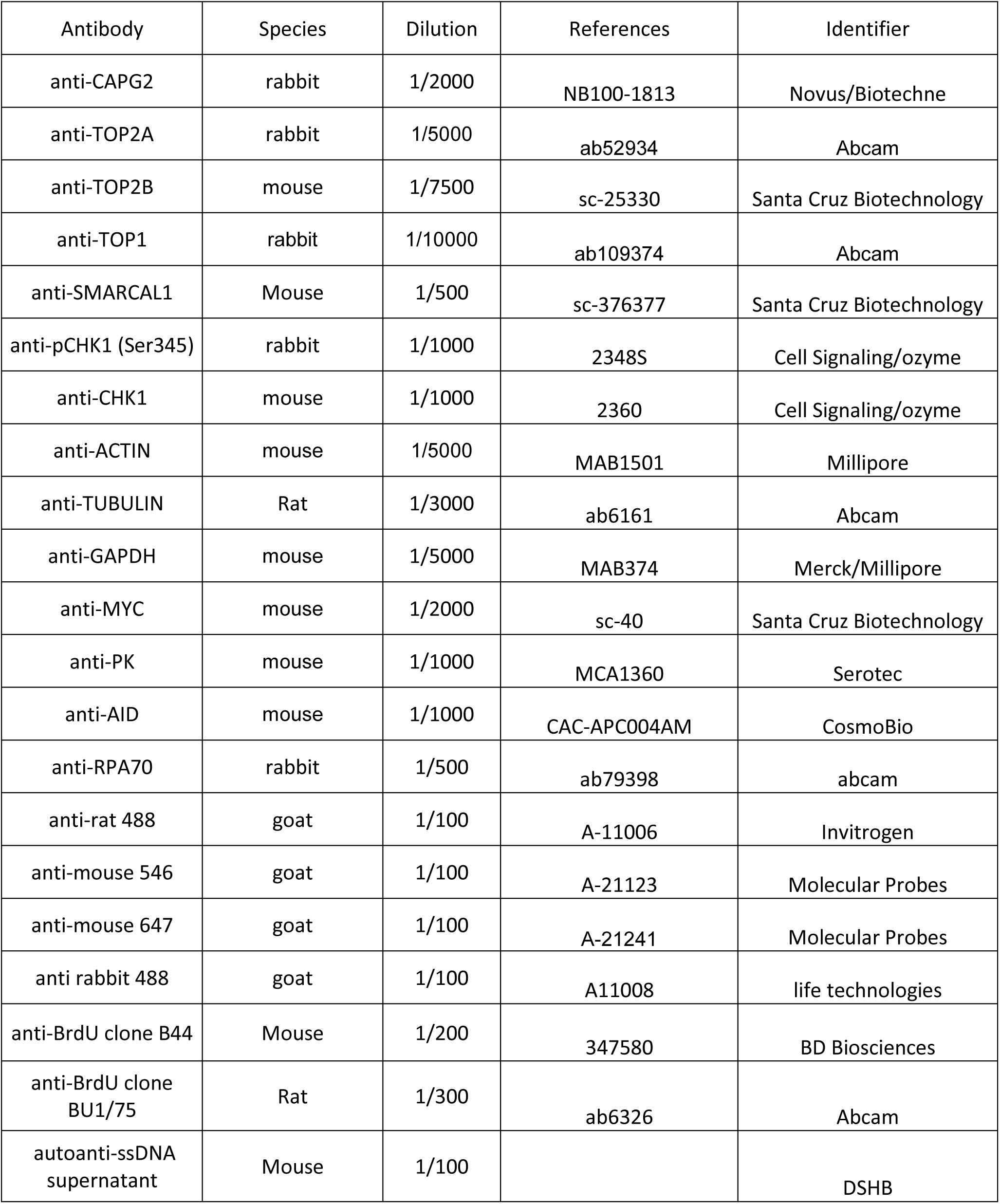
Antibodies used in this study.

Images were acquired by using Zeiss Apotome microscope. The length of tracks was measured by using ImageJ software and analysis was performed using GraphPad Prism analysis. The lengths of at least 100 CldU/IdU tracks were measured per sample.

### Electronic Microscopy

For EM analysis of replication intermediates, 4 µg/mL doxycycline was added to the media of actively replicating shCtrl or shCAPG2 Hela cells for 72 hours. In the last 2 hours of doxycycline induction, the cells were treated with either 4 mM hydroxyurea alone or combined 4 mM hydroxyurea with 50 µM mirin. Non-treated samples were also included. Cells were immediately collected after treatment, washed, and DNA was cross-linked by incubating with 10 µg/mL 4,5’,8-trimethylpsoralen (T6137, Millipore Sigma) followed by a 3-minute exposure to 366 nm UV light on a precooled metal block, for a total of three rounds. Cells were lysed and genomic DNA was isolated from the nuclei by proteinase K (25530-015, Life Technologies) digestion and chloroform-isoamyl alcohol extraction. Genomic DNA was precipitated using isopropanol and digested with *Pvu*II HF (R3151S, New England Biolabs) with the appropriate buffer for 4 hours at 37°C. Replication intermediates were enriched on QIAGEN plasmid mini kit columns (12123, QIAGEN) and concentrated using Amicon Ultra size exclusion columns (UFC510096, Millipore Sigma). Samples were prepared for EM visualization by spreading the enriched, concentrated DNA on a carbon-coated grid in the presence of benzyl-dimethyl-alkylammonium chloride (B6295, Millipore Sigma) as well as n,n-dimethylformamide (227056, Millipore Sigma), followed by low angle platinum rotary shadowing. Images were obtained on a JEOL JEM-1400 electron microscope using a bottom mounted AMT XR401 camera. Analysis was performed using ImageJ software (National Institute of Health). EM analysis allows for distinguishing duplex DNA—which is expected to appear as a 10 nm thick fiber after the platinum coating step necessary for EM visualization—from ssDNA, which has a reduced thickness of 5-7 nm and appears lighter in contrast as that of dsDNA. Criteria used for the assignment of a three-way junction, indicative of a replication fork, include the joining of three DNA fibers into a single junction, with two generally symmetrical daughter strands and a single parental strand. Reversed replication forks consist of four DNA fibers joined at a single junction, consisting of two commonly symmetrical daughter strands, one parental strand and the addition of a typically shorter fourth strand, representative of the reversed arm. The length of the two daughter strands corresponding to the newly replicated duplex oftentimes are equal (b=c), whereas the length of the parental arm and the regressed arm can vary (a ≠ b = c ≠ d). Conversely, canonical Holliday junction structures will be characterized by arms of equal length (a = b, c = d). Particular attention is paid to the junction of the reversed replication fork in order to observe the presence of a bubble structure, indicating that the junction is opened up and that it is simply not the result of the occasional crossover of two DNA molecules. These four-way junctions of reversed replication forks may also be collapsed and other indicators such as daughter strand symmetry, presence of single-stranded DNA at the junction or the entire structure itself, all are considered during analysis^76^. The frequency of reversed forks in a sample is computed using the Prism software.

### Cell survival assay

200 cells were platted and treated during 24 hours with HU concentration as indicated. Then, cells were washed and grown during one week. The cells were fixed with cold 80% methanol for 10 min and stained with crystal violet. Distinct colonies were counted.

For drug sensitivity assay, 200 cells were plated. The next day, HU was added as indicated and cells were grown during 3 days. Cells were incubated with Cell Proliferation Reagent WST1 for 1 hours and the cell colorimetry was read with an ELISA reader.

### Immunofluorescence microscopy

Cells were incubated 5min at 4°C with CSK buffer (20 mM HEPES, 50 mM NaCl, 300 mM sucrose, 3 mM MgCl2, 0.5% Triton X-100, one tablet of protease EDTA-free inhibitor). Then, cells were fixed with 2% paraformaldehyde in PBS for 10min and permeabilized with 0.5% Triton X-100 for 10 min. The coverslips were incubated 1 hour with an anti-RPA70 antibody (1/500) at 37°C and then with a secondary antibody conjugated to an Alexa Fluor dye (1/500) for 1 hour at 37°C. The click-it reaction (PBS, 3mM CuSO4, 29 μM A647 azide dye, 10 mM sodium ascorbate) was performed for 30min at room temperature in the dark, followed by a DAPI staining and ProlongGold mouting. Images were acquired by using Zeiss Apotome microscope and analysed with CellProfiler software.

### iPOND

iPOND data was extracted from^52^.

### Statistical analysis

All numbers of biological replicates are listed in the figure legends. To compare statistical significance between two distributions, the Mann-Whitney rank-sum test was used. All statistical tests were performed with GraphPad Prism 7.

## Data availability

The data sets generated and/or analyzed during the current study are available from the corresponding authors on reasonable request. The NGS data sets generated and analyzed during the current study are available in the GEO repository, accession number: GSE280436.

## ACKNOWLEDGEMENTS

We thank Benjamin Pardo and other members of the Pasero laboratory for discussions and comments on the manuscript. We also thank Carol Featherstone of Plume Scientific Communication Services for editorial assistance during the preparation of the manuscript. We are grateful to the DNA combing facility of Montpellier (BioCampus) for silanized coverslips. We thank the imaging facility MRI, member of the national infrastructure France-BioImaging supported by the French National Research Agency (ANR-10-INBS-04, «Investments for the future»). MD thanks the French MRES for fellowship. MD also thanks the Fondation pour la Recherche Médicale (FRM) for support (FDT202304016554). This work was supported by grants to AL from the Agence Nationale pour la Recherche (ANR-20-CE12-0016-01) and Fondation ARC (ARCPJA2023060006648). Work in the PP lab is also supported by grants from the Institut National du Cancer (INCa), the Ligue Nationale Contre le Cancer (équipe labéllisée) and the Fondation MSDAvenir. Work in the A.V. lab is supported by the National Cancer Institute (NCI) grants R01CA237263 and R01CA248526, and by the Alvin J. Siteman Cancer Center Siteman Investment Program (supported by The Foundation for Barnes-Jewish Hospital, Cancer Frontier Fund).

## AUTHOR CONTRIBUTIONS

Conceptualization, MD, AD, YLL, PP and AL; Methodology, MD, AD, AB, ATV, JJ and AL; Supervision: AV, YLL, PP and AL; Project Administration, PP and AL; Funding Acquisition, AV, PP and AL; Writing – Original Draft, MD, PP and AL; Writing – Review & Editing, MD, JJ, AV, YLL, PP and AL

## REFERENCES

1 Pasero, P. & Vindigni, A. Nucleases acting at stalled forks: how to reboot the replication program with a few shortcuts. Annual Review in Genetics 51, 477–499, (2017).

2 Neelsen, K. J. & Lopes, M. Replication fork reversal in eukaryotes: from dead end to dynamic response. Nat Rev Mol Cell Biol 16, 207–220, (2015).

3 Adolph, M. B. & Cortez, D. Mechanisms and regulation of replication fork reversal. DNA Repair 141, 103731, (2024).

4 Quinet, A., Carvajal-Maldonado, D., Lemacon, D. & Vindigni, A. DNA Fiber Analysis: Mind the Gap! Methods Enzymol 591, 55–82, (2017).

5 Postow, L. et al. Positive torsional strain causes the formation of a four-way junction at replication forks. J Biol Chem 276, 2790–2796, (2001).

6 Fierro-Fernandez, M., Hernandez, P., Krimer, D. B., Stasiak, A. & Schvartzman, J. B. Topological locking restrains replication fork reversal. PNAS 104, 1500–1505, (2007).

7 Tian, T. et al. The ZATT-TOP2A-PICH Axis Drives Extensive Replication Fork Reversal to Promote Genome Stability. Mol Cell 81, 198–211.e196, (2021).

8 Pommier, Y., Nussenzweig, A., Takeda, S. & Austin, C. Human topoisomerases and their roles in genome stability and organization. Nat Rev Mol Cell Biol 23, 407–427, (2022).

9 Hirano, T. Condensin-Based Chromosome Organization from Bacteria to Vertebrates. Cell 164, 847–857, (2016).

10 Ono, T. et al. Differential contributions of condensin I and condensin II to mitotic chromosome architecture in vertebrate cells. Cell 115, 109–121, (2003).

11 Freeman, L., Aragon-Alcaide, L. & Strunnikov, A. The condensin complex governs chromosome condensation and mitotic transmission of rDNA. J Cell Biol 149, 811–824, (2000).

12 D’Ambrosio, C. et al. Identification of cis-acting sites for condensin loading onto budding yeast chromosomes. Genes Dev 22, 2215–2227, (2008).

13 Doughty, T. W., Arsenault, H. E. & Benanti, J. A. Levels of Ycg1 Limit Condensin Function during the Cell Cycle. PLoS Genet 12, e1006216, (2016).

14 Leonard, J. et al. Condensin Relocalization from Centromeres to Chromosome Arms Promotes Top2 Recruitment during Anaphase. Cell Rep 13, 2336–2344, (2015).

15 Thadani, R., Kamenz, J., Heeger, S., Muñoz, S. & Uhlmann, F. Cell-Cycle Regulation of Dynamic Chromosome Association of the Condensin Complex. Cell Rep 23, 2308–2317, (2018).

16 Dinda, M. et al. Fob1-dependent condensin recruitment and loop extrusion on yeast chromosome III. PLoS Genet 19, e1010705, (2023).

17 Sakamoto, T. et al. Condensin II alleviates DNA damage and is essential for tolerance of boron overload stress in Arabidopsis. Plant Cell 23, 3533–3546, (2011).

18 Ono, T., Yamashita, D. & Hirano, T. Condensin II initiates sister chromatid resolution during S phase. J Cell Biol 200, 429–441, (2013).

19 Fazzio, T. G. & Panning, B. Condensin complexes regulate mitotic progression and interphase chromatin structure in embryonic stem cells. J Cell Biol 188, 491–503, (2010).

20 Woodward, J. et al. Condensin II mutation causes T-cell lymphoma through tissue-specific genome instability. Genes Dev 30, 2173–2186, (2016).

21 Kakui, Y. et al. Fission yeast condensin contributes to interphase chromatin organization and prevents transcription-coupled DNA damage. Genome Biol 21, 272, (2020).

22 Xu, X., Nakazawa, N. & Yanagida, M. Condensin HEAT subunits required for DNA repair, kinetochore/centromere function and ploidy maintenance in fission yeast. PLoS One 10, e0119347, (2015).

23 D’Ambrosio, C. et al. Identification of cis-acting sites for condensin loading onto budding yeast chromosomes. Genes & Development 22, 2215–2227, (2008).

24 Poli, J. et al. dNTP pools determine fork progression and origin usage under replication stress. EMBO J 31, 883–894, (2012).

25 Yabuki, N., Terashima, H. & Kitada, K. Mapping of early firing origins on a replication profile of budding yeast. Genes Cells 7, 781–789, (2002).

26 Crabbé, L. et al. Analysis of replication profiles reveals key role of RFC-Ctf18 in yeast replication stress response. Nature Structural & Molecular Biology 17, 1391–1397, (2010).

27 Delamarre, A. et al. MRX Increases Chromatin Accessibility at Stalled Replication Forks to Promote Nascent DNA Resection and Cohesin Loading. Mol Cell 77, 395–410.e393, (2020).

28 Aono, N., Sutani, T., Tomonaga, T., Mochida, S. & Yanagida, M. Cnd2 has dual roles in mitotic condensation and interphase. Nature 417, 197–202, (2002).

29 Lancaster, L., Patel, H., Kelly, G. & Uhlmann, F. A role for condensin in mediating transcriptional adaptation to environmental stimuli. Life Sci Alliance 4, (2021).

30 Tourrière, H., Saksouk, J., Lengronne, A. & Pasero, P. Single-molecule Analysis of DNA Replication Dynamics in Budding Yeast and Human Cells by DNA Combing. Bio-protocol 7, e2305, (2017).

31 Baxter, J. et al. Positive Supercoiling of Mitotic DNA Drives Decatenation by Topoisomerase II in Eukaryotes. Science 331, 1328–1332, (2011).

32 Charbin, A., Bouchoux, C. & Uhlmann, F. Condensin aids sister chromatid decatenation by topoisomerase II. Nucleic Acids Res 42, 340–348, (2014).

33 Dyson, S., Segura, J., Martínez-García, B., Valdés, A. & Roca, J. Condensin minimizes topoisomerase II-mediated entanglements of DNA in vivo. Embo j 40, e105393, (2021).

34 Martínez-García, B. et al. Condensin pinches a short negatively supercoiled DNA loop during each round of ATP usage. Embo j 42, e111913, (2023).

35 Bermejo, R. et al. Top1- and Top2-mediated topological transitions at replication forks ensure fork progression and stability and prevent DNA damage checkpoint activation. Genes Dev. 21, 1921–1936, (2007).

36 Dungrawala, H. et al. The Replication Checkpoint Prevents Two Types of Fork Collapse without Regulating Replisome Stability. Molecular Cell 59, 998–1010, (2015).

37 Coquel, F. et al. SAMHD1 acts at stalled replication forks to prevent interferon induction Nature 557, 57–61, (2018).

38 Heuzé, J. et al. RNase H2 degrades toxic RNA:DNA hybrids behind stalled forks to promote replication restart. EMBO J, e113104, (2023).

39 Zellweger, R. et al. Rad51-mediated replication fork reversal is a global response to genotoxic treatments in human cells. The Journal of Cell Biology 208, 563–579, (2015).

40 Andrs, M. et al. Excessive reactive oxygen species induce transcription-dependent replication stress. Nat Commun 14, 1791, (2023).

41 Schlacher, K. et al. Double-strand break repair-independent role for BRCA2 in blocking stalled replication fork degradation by MRE11. Cell 145, 529–542, (2011).

42 Vindigni, A. & Lopes, M. Combining electron microscopy with single molecule DNA fiber approaches to study DNA replication dynamics. Biophys Chem 225, 3–9, (2017).

43 Bai, G. et al. HLTF Promotes Fork Reversal, Limiting Replication Stress Resistance and Preventing Multiple Mechanisms of Unrestrained DNA Synthesis. Mol Cell 78, 1237–1251.e1237, (2020).

44 Leung, W. et al. ATR protects ongoing and newly assembled DNA replication forks through distinct mechanisms. Cell Rep 42, 112792, (2023).

45 Qiu, S., Jiang, G., Cao, L. & Huang, J. Replication Fork Reversal and Protection. Front Cell Dev Biol 9, 670392, (2021).

46 Liu, W., Krishnamoorthy, A., Zhao, R. & Cortez, D. Two replication fork remodeling pathways generate nuclease substrates for distinct fork protection factors. Sci Adv 6, eabc3598, (2020).

47 Kimura, K. & Hirano, T. ATP-dependent positive supercoiling of DNA by 13S condensin: a biochemical implication for chromosome condensation. Cell 90, 625–634, (1997).

48 Manders, E. M., Kimura, H. & Cook, P. R. Direct imaging of DNA in living cells reveals the dynamics of chromosome formation. J Cell Biol 144, 813–821, (1999).

49 Akai, Y. et al. Opposing role of condensin hinge against replication protein A in mitosis and interphase through promoting DNA annealing. Open Biol 1, 110023, (2011).

50 Aze, A., Sannino, V., Soffientini, P., Bachi, A. & Costanzo, V. Centromeric DNA replication reconstitution reveals DNA loops and ATR checkpoint suppression. Nature Cell Biology 18, 684, (2016).

51 Sutani, T. et al. Condensin targets and reduces unwound DNA structures associated with transcription in mitotic chromosome condensation. Nat Commun 6, 7815, (2015).

52 Lebdy, R. et al. The nucleolar protein GNL3 prevents resection of stalled replication forks. EMBO Rep 24, e57585, (2023).

53 Wessel, S. R., Mohni, K. N., Luzwick, J. W., Dungrawala, H. & Cortez, D. Functional Analysis of the Replication Fork Proteome Identifies BET Proteins as PCNA Regulators. Cell Rep 28, 3497–3509.e3494, (2019).

54 Iwasaki, O. et al. Interaction between TBP and Condensin Drives the Organization and Faithful Segregation of Mitotic Chromosomes. Mol Cell 59, 755–767, (2015).

55 Toselli-Mollereau, E. et al. Nucleosome eviction in mitosis assists condensin loading and chromosome condensation. Embo j 35, 1565–1581, (2016).

56 Haase, J. et al. The TFIIH complex is required to establish and maintain mitotic chromosome structure. Elife 11, (2022).

57 Colin, L. et al. Condensin positioning at telomeres by shelterin proteins drives sister-telomere disjunction in anaphase. Elife 12, (2023).

58 Pradhan, B. et al. SMC complexes can traverse physical roadblocks bigger than their ring size. Cell Rep 41, 111491, (2022).

59 Guérin, T. M. et al. Condensin-Mediated Chromosome Folding and Internal Telomeres Drive Dicentric Severing by Cytokinesis. Mol Cell 75, 131–144.e133, (2019).

60 Analikwu, B. T. et al. Telomere protein arrays stall DNA loop extrusion by condensin. bioRxiv, 2023.2010.2029.564563, (2023).

61 Mariezcurrena, A. & Uhlmann, F. Observation of DNA intertwining along authentic budding yeast chromosomes. Genes Dev 31, 2151–2161, (2017).

62 Orlandini, E., Marenduzzo, D. & Michieletto, D. Synergy of topoisomerase and structural-maintenance-of-chromosomes proteins creates a universal pathway to simplify genome topology. Proc Natl Acad Sci U S A 116, 8149–8154, (2019).

63 Elbatsh, A. M. O. et al. Distinct Roles for Condensin’s Two ATPase Sites in Chromosome Condensation. Mol Cell 76, 724–737.e725, (2019).

64 Samejima, K. et al. Mitotic chromosomes are compacted laterally by KIF4 and condensin and axially by topoisomerase IIα. J Cell Biol 199, 755–770, (2012).

65 Sen, N. et al. Physical Proximity of Sister Chromatids Promotes Top2-Dependent Intertwining. Mol Cell 64, 134–147, (2016).

66 Piskadlo, E., Tavares, A. & Oliveira, R. A. Metaphase chromosome structure is dynamically maintained by condensin I-directed DNA (de)catenation. Elife 6, (2017).

67 Eeftens, J. M. et al. Real-time detection of condensin-driven DNA compaction reveals a multistep binding mechanism. Embo j 36, 3448–3457, (2017).

68 Kim, E., Gonzalez, A. M., Pradhan, B., van der Torre, J. & Dekker, C. Condensin-driven loop extrusion on supercoiled DNA. Nat Struct Mol Biol 29, 719–727, (2022).

69 Liu, W. et al. RAD51 bypasses the CMG helicase to promote replication fork reversal. Science 380, 382–387, (2023).

70 Bazett-Jones, D. P., Kimura, K. & Hirano, T. Efficient supercoiling of DNA by a single condensin complex as revealed by electron spectroscopic imaging. Mol Cell 9, 1183–1190, (2002).

71 Tittel-Elmer, M. et al. Cohesin Association to Replication Sites Depends on Rad50 and Promotes Fork Restart. Molecular Cell 48, 98–108, (2012).

72 Jeppsson, K. et al. The chromosomal association of the Smc5/6 complex depends on cohesion and predicts the level of sister chromatid entanglement. PLoS Genet 10, e1004680, (2014).

73 Bonner, J. N. & Zhao, X. Replication-Associated Recombinational Repair: Lessons from Budding Yeast. Genes (Basel) 7, (2016).

74 Meng, X. & Zhao, X. Replication fork regression and its regulation. FEMS Yeast Research 17, fow110-fow110, (2017).

75 Lin, Y. L. et al. Feline immunodeficiency virus vectors for efficient transduction of primary human synoviocytes: application to an original model of rheumatoid arthritis. Hum Gene Ther 15, 588–596, (2004).

76 Neelsen, K. J., Chaudhuri, A. R., Follonier, C., Herrador, R. & Lopes, M. Visualization and interpretation of eukaryotic DNA replication intermediates in vivo by electron microscopy. Methods Mol Biol 1094, 177–208, (2014).

77 Ribeyre, C. et al. Nascent DNA Proteomics Reveals a Chromatin Remodeler Required for Topoisomerase I Loading at Replication Forks. Cell Rep 15, 300–309, (2016).

