## Supplementary figures for "Condensin and topoisomerases cooperate to relieve topological stress at stalled replication forks"

**Supplementary Fig. 1 Condensin is recruited to replication stress sites in budding yeast in a Rad50-dependent manner. a** Condensin promotes cell growth in the presence of MMS. *SMC4*-AID degron cells complemented or not with WT *SMC4* were grown for 4 days on YPD medium containing 0.033% MMS and a 0 to 25 µM gradient of auxin (IAA). **b** Condensin is enriched at active origins in HU-arrested cells. Genome-wide distribution of Brn1-$PK_9$ in wild type (untagged), Brn1-$PK_9$ and *rad50D* Brn1-$PK_9$ cells arrested in $G_1$ with a-factor ($G_1$) or released for 60 minutes into S phase in the presence of 200 mM HU. Representative regions on chromosomes III and VI are shown. The positions of the early origins *ARS306*, *ARS606* and *ARS607* are indicated. **c** Condensin enrichment at stressed forks depends on the MRX subunit Rad50. WT and *rad50D* cells expressing *BRN1-PK$_9$* were released for 60 minutes from $G_1$ into S phase in the presence of 200 mM HU. Brn1 enrichment was determined by ChIP-qPCR at the indicated distances to *ARS305* and *ARS306*. Brn1 enrichment was normalized to unreplicated regions. Mean and SD correspond to three independent experiments.

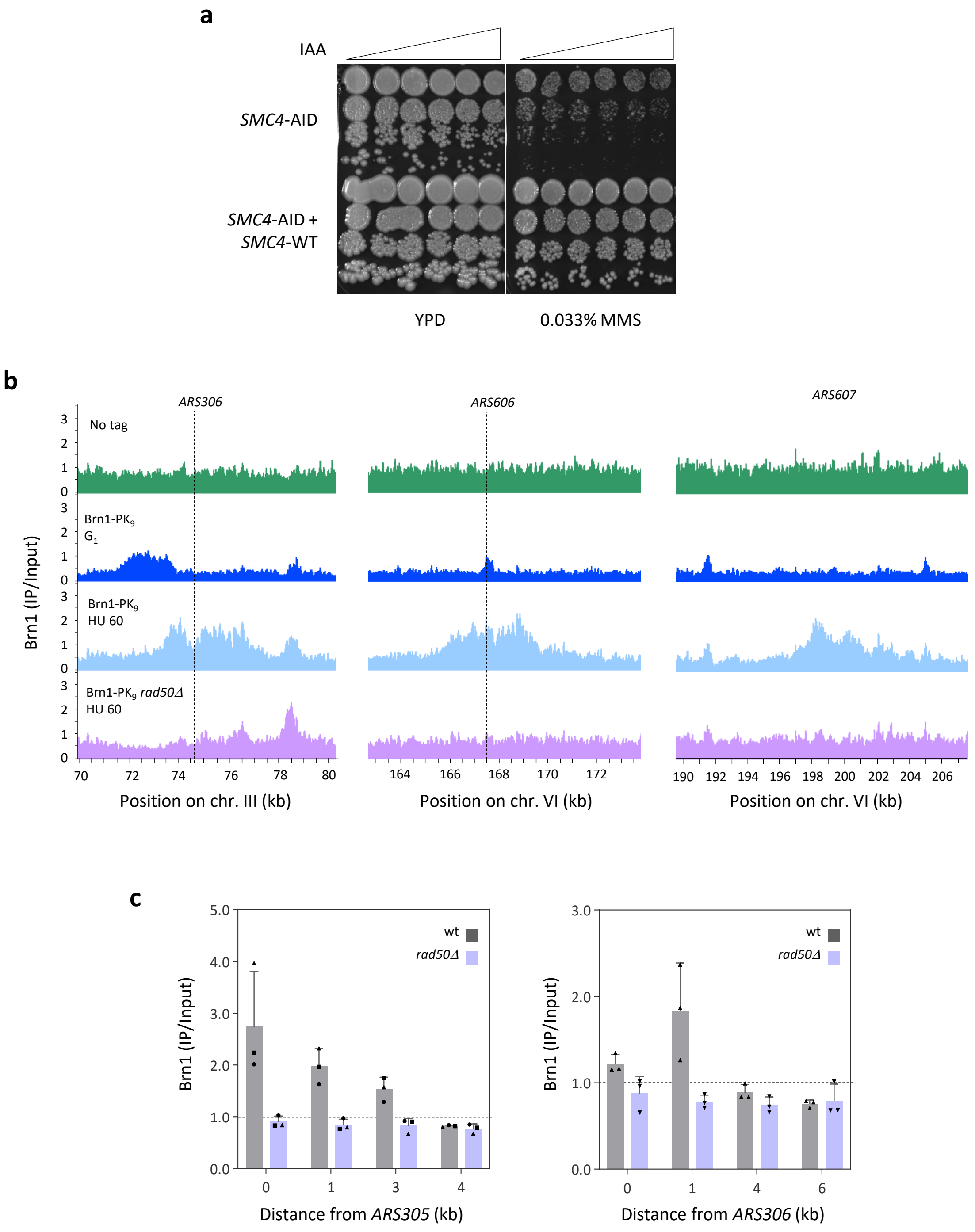


Supplementary Figure 1

**Supplementary Fig. 2 Condensin is dispensable for normal fork progression and for checkpoint activation, but contributes to fork restart in budding yeast. a, b** The replication checkpoint is functional in cells depleted for the condensin subunit Ycg1. WT and pMET-*YCG1*-$PK_3$-AID cells grown in medium without methionine were synchronized in $G_1$ with α-factor and released into S phase in the presence of 200 mM HU. Cells were shifted to YPD medium after 60 minutes of $\alpha$-factor addition and auxin was added 100 minutes before $G_1$ release to deplete Ycg1. Origin firing was monitored by DNA copy number variation. DNA was extracted and analyzed by qPCR with primers corresponding to the indicated early and late origins. DNA content was normalized to an unreplicated region in two independent experiments. The repression of the late origins *ARS809*, *ARS911* and *ARS1212* is indicative of the timely activation of the replication checkpoint in the presence of HU. **c** Fork progression is impaired in HU-treated *smc2-8* mutants. A representative experiment from Fig. 2a (n=3) is shown. ****: p < 0.0001; ns: non-significant, Mann–Whitney rank-sum test. **d** Ycg1-depleted cells show a slower fork progression in the presence of 200 mM HU. Cells were grown as described in panel a. BrdU was added after 60 minutes in HU for 120 minutes. ****: p < 0.0001, Mann–Whitney rank-sum test. **e** Condensin is dispensable for normal fork progression. Exponentially growing *YCG1*-$PK_3$ and pMET-*YCG1*-$PK_3$-AID cells grown in medium lacking methionine were transferred for 30 minutes into YPD medium before auxin addition to deplete Ycg1. EdU was then added for 15 minutes to label newly replicated DNA and the length of EdU tracks was measured by DNA combing. Box and whiskers indicate 25th–75th and 10th–90th percentiles (n=3). **f** Condensin is required for timely fork restart after MMS exposure. A representative experiment from Fig. 2b (n=3) is shown. ****: p < 0.0001; ns: non-significant, Mann–Whitney rank-sum test. **g** Top1 depletion restores fork restart in the absence of condensin. A representative experiment from Fig. 2d (n=3) is shown. ****: p < 0.0001; ns: non-significant, Mann–Whitney rank-sum test. **h** Condensin acts with Top2 to promote fork restart after MMS exposure. A representative experiment from Fig. 2e (n=3) is shown. ****: p < 0.0001; ns: non-significant, Mann–Whitney rank-sum test. **i, j** Western blot analysis of Top1 and Top2 depletion in the experiments described in panels g and h.

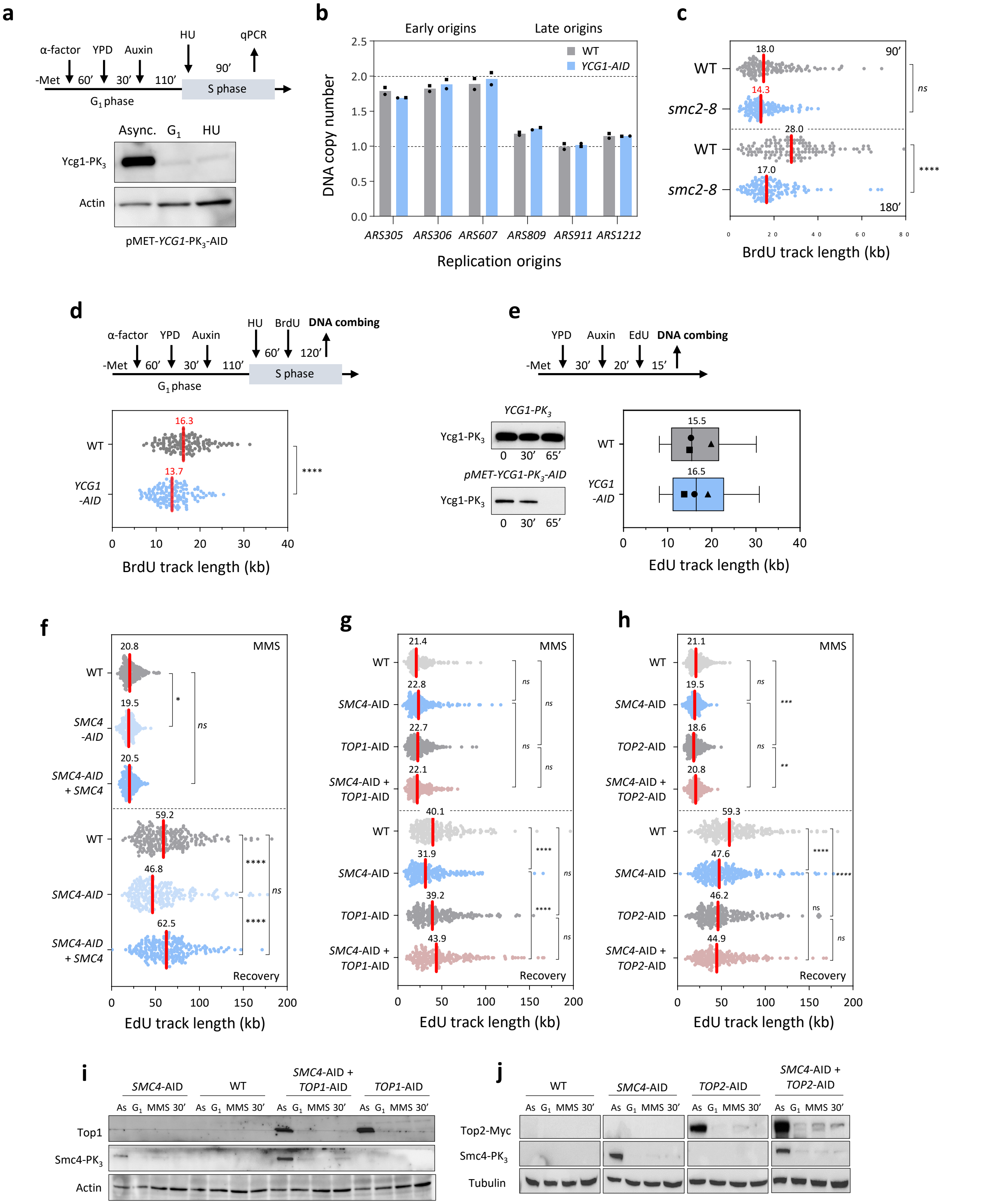


Supplementary Figure 2

**Supplementary Fig. 3 Condensin promotes cell growth under replication stress conditions. a** Condensin I and II are dispensable for normal fork progression. A representative experiment from Fig. 3b is shown. *: $p < 0.1$; ns: non-significant, Mann–Whitney rank-sum test. **b** Condensin II is required for growth in the presence of HU. HeLa-S3 cells were transfected with siRNAs (siCtrl or siCAPG2) for 48 hours. Cells were incubated with increasing doses of HU for 3 days. Cell proliferation was quantified and normalized to control cells at day 0. Mean and SD are indicated (n=3). **c** Condensin II promotes cell survival after acute HU exposure. HeLa-S3 cells were transfected with siRNAs (siCtrl or siCAPG2) for 48 hours. Cells were incubated for 24 hours with 1 or 2 mM of HU. Colonies were counted after 7 days and the percentage of survival for each cell line was normalized to untreated condition. Mean and SD are indicated (n = 3). **d** Condensin II is required for fork slowing after exposure to a low dose of HU. A representative experiment from Fig. 3e is shown. ****: $p < 0.0001$; ns: non-significant, Mann–Whitney rank-sum test. **e** Condensin II, but not Condensin I, promotes the resection of nascent DNA at HU-arrested forks. HeLa-S3 cells were transfected with siCtrl, siCAPG or siCAPG2 for 48 hours and were sequentially labeled for 15 minutes with IdU and CldU. Cells were either collected immediately or treated for 2 hours with 4 mM HU before DNA fiber analysis. The ratio of CldU to IdU track length is shown (n=3). ****: $p < 0.0001$; *: $p < 0.1$, Mann–Whitney rank-sum test. **f** Condensin II promotes the resection of nascent DNA at HU-arrested forks in U2OS cells. U2OS cells were transfected with siCtrl and siCAPG2 for 48 h. After sequential labelling of IdU and CldU for 15 min, cells were either collected immediately or treated for 2 hours with 4 mM HU before DNA fiber analysis. The ratio of CldU to IdU track length was plotted for two independent experiments. Box and whiskers indicate median, 25th–75th and 10th–90th percentiles.

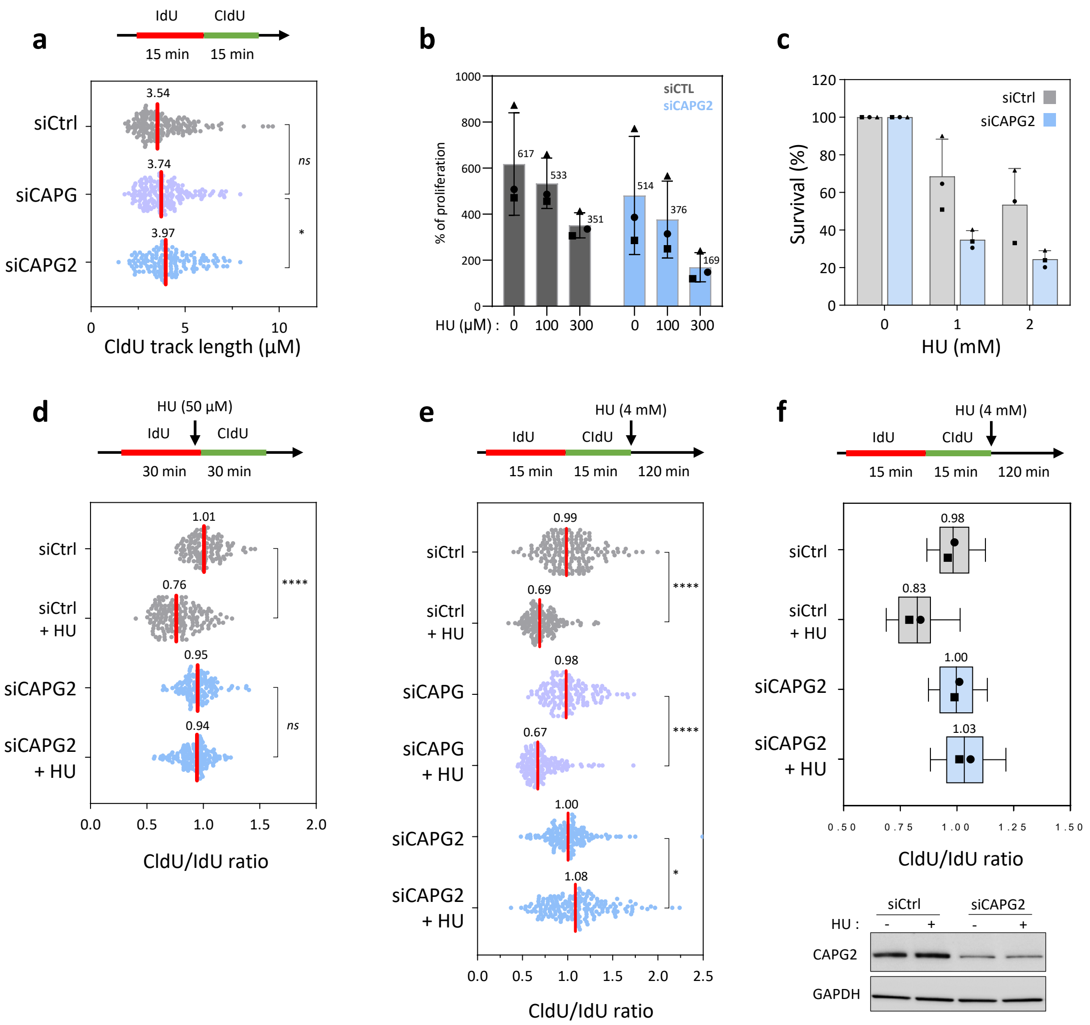


Supplementary Figure 3

**Supplementary Fig. 4 Condensin II acts with DNA translocases to promote fork reversal. a** Control (shCtrl) and CAPG2-depleted cells (shCAPG2) HeLa-S3 cells were treated for 72 hours with 10 μM/ml doxycycline and then for 2 hours with 4 mM HU, under the same experimental conditions used for EM analysis (Fig. 4a and b). Fork resection was monitored by DNA fiber spreading after sequential labelling with IdU and CldU for 15 minutes. Cells were either collected immediately or treated for 2 hours with 4 mM HU before DNA fiber analysis. Box and whiskers indicate median, 25th–75th and 10th–90th percentiles (n=3). **b** CAPG2 depletion was analyzed by Western blotting. CHK1 phosphorylation was detected with an anti-p-CHK1 (S345) antibody. Actin was used as loading control. **c** Defective fork slowdown in SMARCAL1-deficient cells is rescued by CAPG2 depletion. A representative experiment from Fig. 4d is shown. ****: $p < 0.0001$; *: $p < 0.1$; Mann–Whitney rank-sum test. **d** Analysis of CAPG2 and SMARCAL1 depletion by Western blotting. GAPDH is used as loading control. **e** CAPG2 depletion restores fork resection in SMARCAL1-depleted cells. A representative experiment from Fig. 4e is shown. ****: $p < 0.0001$; ns: non-significant; Mann–Whitney rank-sum test. **f** Analysis of CAPG2 and SMARCAL1 depletion by Western blotting. CHK1 phosphorylation was detected with an anti-pCHK1 (S345) antibody. Total CHK1 was used as loading control. **g** CAPG2 depletion restores fork slowdown in both SMARCAL1- and FBH1-depleted cells. U2OS cells were transfected for 48 hours with siCtrl, siCAPG2, siFBH1, or co-transfected with siSMARCAL1 as indicated. Cells were first labelled for 30 minutes with IdU, and CldU was added for another 30 minutes in the presence of 50 μM HU. The ratio of CldU to IdU track length was plotted for four independent experiments. Box and whiskers indicate median, 25th–75th and 10th–90th percentiles.

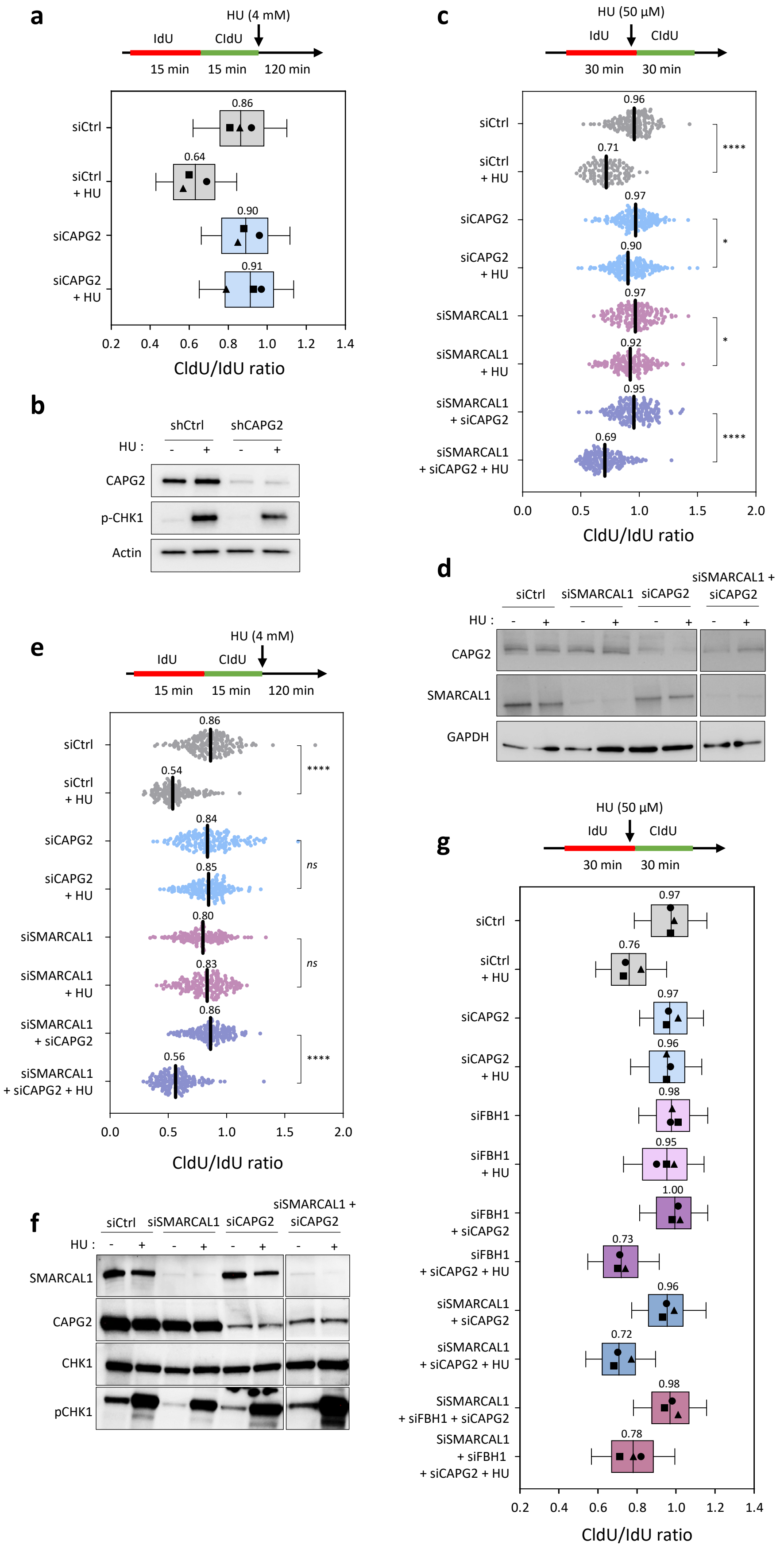


Supplementary Figure 4

**Supplementary Fig. 5 Interplay between condensin II and topoisomerases at stalled replication forks. a** Condensin II acts with TOP2A to promote fork slowing. A representative experiment from Fig. 5a is shown. ****: $p < 0.0001$; ***: $p < 0.001$; ns: non-significant; Mann–Whitney rank-sum test. **b** TOP2A and CAPG2 levels were analyzed by Western blotting using GAPDH as loading control. **c** Condensin II acts with TOP2A to promote fork resection. A representative experiment from Fig. 5b is shown. ****: $p < 0.0001$; **: $p < 0.01$; ns: non-significant; Mann–Whitney rank-sum test. **d** Western blot analysis of TOP2A and CAPG2 depletion. CHK1 activation was detected with an anti-pCHK1 (S345) antibody. Total CHK1 and actin were used as loading controls. **e** TOP2B act with CAPG2 to promote fork resection. HeLa-S3 cells were transfected with siCtrl, siCAPG2, siTOP2B or co-transfected with siTOP2B and siCAPG2 for 48 h. After sequential labelling of IdU and CldU for 15 minutes, cells were either collected immediately or treated for 2 hours with 4 mM HU before DNA fiber analysis. The ratio of CldU to IdU track length is shown for three independent experiments. Box and whiskers correspond to median, 25th–75th and 10th–90th percentiles. Median length is indicated. **f** Western blot analysis of TOP2B and CAPG2 levels. CHK1 activation was detected with an anti-pCHK1 (S345) antibody. GAPDH was used as loading control. **g** TOP1 depletion restores fork slowing in CAPG2-deficient cells. A representative experiment from Fig. 5c is shown. ****: $p < 0.0001$; *: $p < 0.1$; ns: non-significant; Mann–Whitney rank-sum test. **h** TOP1 and CAPG2 levels were analyzed by Western blotting using total CHK1 is used as loading control. **i** TOP1 depletion restores fork resection in CAPG2-deficient cells. A representative experiment from Fig. 5d is shown. ****: $p < 0.0001$; ns: non-significant; Mann–Whitney rank-sum test. **j** Western blot analysis of TOP1 and CAPG2 levels and CHK1 activation. Total CHK1 and actin were used as loading controls. **k** TOP1 depletion restores fork resection in SMARCAL1-deficient cells. A representative experiment from Fig. 5e is shown. ****: $p < 0.0001$; ns: non-significant; Mann–Whitney rank-sum test. **l** TOP1 depletion restores fork slowing in SMARCAL1-deficient cells. HeLa-S3 cells were transfected with siCtrl, siSMARCAL1, siTOP1, or co-transfected with siSMARCAL1 and siTOP1 for 48 h. Cells were labeled and analyzed by DNA fiber spreading as described in Fig. 5b (n=2). **m** Depletion of TOP2A does not restore fork resection in SMARCAL1-deficient cells. A representative experiment from Fig. 5f is shown. ****: $p < 0.0001$; *: $p < 0.1$; ns: non-significant; Mann–Whitney rank-sum test.

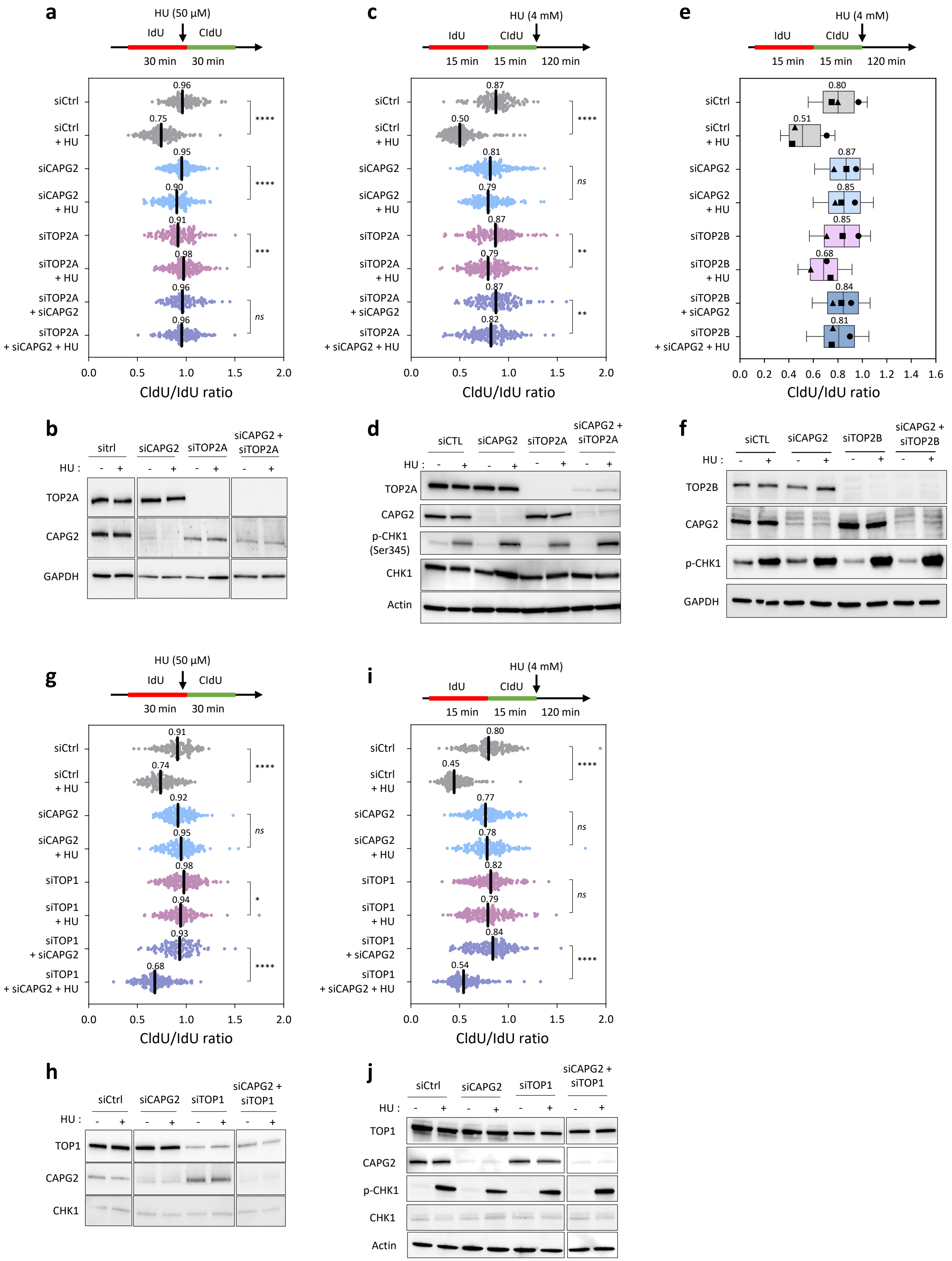


Supplementary Figure 5

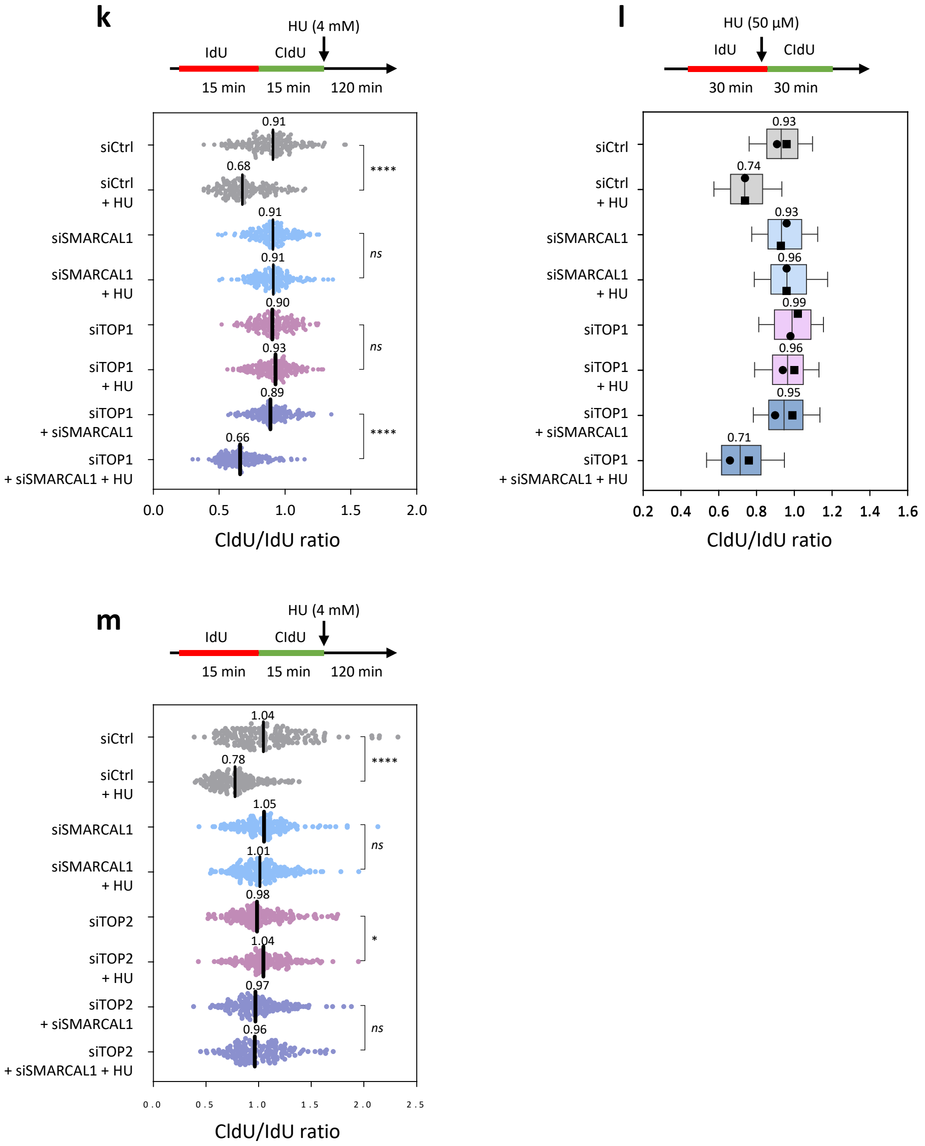


Supplementary Figure 5 (continued)

**Supplementary Fig. 6 RPA-coated ssDNA accumulates at HU-arrested forks in the absence of condensin and Top1 in budding yeast**. **a** WT and pMET-*YCG1*-$PK_3$-AID cells were synchronized in $G_1$ with α-factor and released into S phase in the presence of 200 mM HU. RPA enrichment was determined by ChIP-qPCR at the indicated distances from *ARS607* after normalization to unreplicated loci (n=2). Changes in DNA copy number indicate that forks progress for up to 3 kb from *ARS607* under these conditions. **b** WT, *SMC4*-AID, *top1Δ* and *top1Δ SMC4*-AID cells were arrested in $G_1$ with α-factor and SMC4 was depleted by addition of Auxin for 60 minutes. Then, cells were released into S phase in the presence of 200 mM HU for another 60 minutes. RPA enrichment at indicated distances from *ARS305* was determined by ChIP-qPCR after normalization to unreplicated regions. Mean and SD correspond to three independent experiments.

**a**

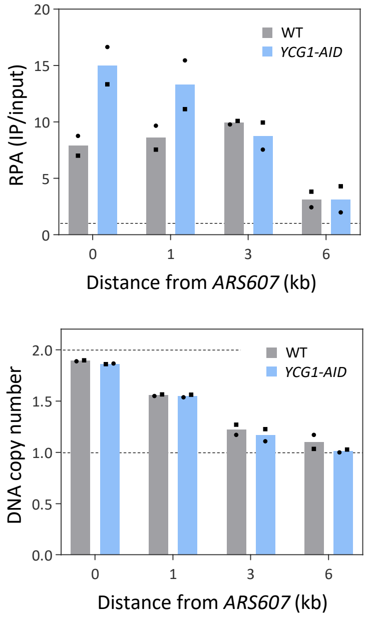


**b**

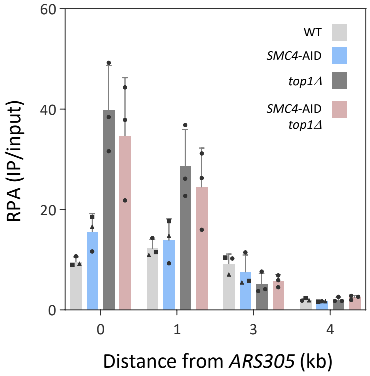


Supplementary Figure 6
